# Vicarious trial-and-error is enhanced during deliberation in human virtual navigation in a translational neuroeconomic task

**DOI:** 10.1101/2020.02.21.954230

**Authors:** Thach Huynh, Keanan Alstatt, Samantha Abram, Neil Schmitzer-Torbert

## Abstract

Foraging tasks can provide valuable insights into decision-making, as animals choose how to allocate limited resources (such as time). In the “Restaurant Row” task, rodents move between several sites to obtain food rewards available after a variable delay, while in a translational version (the “Web-Surf”) lacking the navigation component, humans are offered short videos. Both tasks have provided novel insights into decision-making and have been applied to addiction vulnerability and the impact of drug exposure on decision-making. We tested new tasks (the “Movie Row” and “Candy Row”) which use virtual navigation to determine if the behavioral correlates of human decision-making are more broadly similar to those of rodents, and explored the relationship of task performance to smoking and obesity. Humans navigated a virtual maze presented on standard computers to obtain rewards (either short videos or candy) available after a variable delay. Behavior on the tasks replicated previous results for the Restaurant Row and Web-Surf. In conditions promoting deliberation, decision latency was elevated along with measures of vicarious trial-and-error (VTE), supporting VTE as a shared behavioral index of deliberation across species. Smoking status was not well-related to performance, while high BMI (≥ 25) individuals showed reduced sensitivity to a sunk-costs measure and stronger sensitivity to offer value for offers below the preferred delay. These data support the Movie and Candy Row as translational tools to study decision-making during foraging in humans, providing convergent results with a rodent navigation task and demonstrating the potential to provide novel insights relevant to public health.

---

Decision-making is a key capability, and in some sense a fundamental goal of the nervous system. Rather than a unitary construct, decisions rely on differentiable brain systems, each with a separate computational goal (Redish, 2013). Numerous disadvantageous behaviors can be conceptualized as failures in one or more decision-making systems, and the ability to characterize different facets of the decision-making process is a critical part of the effort to understand normal and pathological decisions.

A number of studies have identified decision making characterized by impulsivity as a risk factor for a range of negative outcomes. Delay-discounting tasks, which are commonly used to assess impulsivity, assess the degree to which rewards available in the future lose value compared to rewards available in the present; these tasks measure impulsivity by presenting a forced-choice between real or hypothetical rewards, while varying the delays associated with each option, and the magnitude of the rewards. Discounting rates are elevated among individuals who smoke, use drugs, have problematic gambling, or are obese (Emery & Levine, 2017; MacKillop et al., 2011). Steep discounting of future rewards may represent a general risk factor that predisposes individuals to be risky behaviors (like drug use), and/or discounting rates may be increased through experiences such as exposure to drugs or engaging in risky behaviors (Naude, Dongelmans, & Faure, 2015).

While discounting of future rewards has been implicated in a range of outcomes involving disadvantageous decision-making, decision-making involves a range of systems (Redish, 2013), and a more complete understanding of any particular condition will require assessment of a wider range of decision-making processes. For example, while both smokers and problematic gamblers show stronger discounting of future rewards, performance during foraging tasks shows a dissociation between these groups, with smokers showing a reduction in exploratory choices (in a multi-armed bandit task, Addicott, Pearson, Wilson, Platt, & McClernon, 2013), while gambling frequency is associated with an increase in exploratory choices (Addicott, Pearson, Kaiser, Platt, & McClernon, 2015). Frequent gambling was also associated with early, suboptimal, exits in a patchy foraging task, also indicating that frequent gamblers showed increased exploration and reduced exploitation.

To better characterize the decision-making process, foraging tasks (in which animals must allocate limited time or effort in order to obtain resources/rewards) are particularly valuable (Stephens, 2008). In one recently described foraging task (Restaurant Row, Steiner, Adam P. & Redish, 2014), rodents navigate on a square track between sites that deliver different flavors of food rewards, which are available after a variable delay. The task was first used to describe behavioral and neural correlates of regret in rats. Rats also demonstrated a behavioral measure of deliberation, vicarious-trial-and-error (VTE), that was sensitive to value: VTE was elevated for difficult decisions, when the delay required to retrieve a reward was close to a rat’s threshold to stay versus skip. Since its original description, the Restaurant Row has also been used with rats and mice to characterize the behavioral correlates of regret (Steiner, Adam P. & Redish, 2014; Sweis, Thomas, & Redish, 2018) and sensitivity to sunk-costs (Sweis et al., 2018).

Restaurant Row has also been used to demonstrate the differential impact exposure to addictive drugs on decision-making. For example, long-term withdrawal from cocaine and heroin in mice produces dissociable effects in mice on Restaurant Row performance (Sweis, Redish, & Thomas, 2018): cocaine-abstinent mice show changes in deliberation, while morphine-abstinent mice show a reduced willingness to quit after initially accepting an offer. And, long-term depression of infralimbic glutamatergic inputs to the nucleus accumbens shell specifically reduces the likelihood that mice will quit after initially accepting an offer, similar to the effects of morphine abstinence (Sweis, Larson, Redish, & Thomas, 2018).

A translational human version of the task, the Web-Surf (Abram, Breton, Schmidt, Redish, & MacDonald III, 2016), uses short (4 second) video clips of different types (kittens, bike accidents, landscapes and dancing) as different types of rewards. Rather than navigating physically between reward sites, humans transition between different video galleries by repeatedly pressing a button. Behavior on the Web-Surf Task converges with several important findings from the Restaurant Row Task, and supports the use of these tasks to study decision-making processes shared across rodents and humans. For example, Sweis and colleagues (2018) demonstrated that both humans and rodents demonstrated a sunk-cost effect on these tasks: after initially accepting an offer, the probability of completing the remainder of a delay increased as a function of the amount of time already invested in each species. In another study, Abram and colleagues (2019) have also shown that the time taken by participants to make a decision on the Web-Surf task is elevated for difficult decisions, similar to findings in rodents.

The degree to which the rodent and human versions of these tasks produce consistent results is an important issue for ongoing research, and for the ability of research using rodents to inform our understanding of human decision-making. One advantage of the Restaurant Row task is the richness of the behavioral measures that can be assessed during navigation, such as head movements and body orientations toward reward sites that are associated with VTE. In the Web-Surf task, the latency to make a decision has been the primary behavioral measure used as an index of deliberation However, other behavioral correlates of deliberation in humans during these kinds of experiential navigation tasks have not been described. Several studies have indicated that eye movements in human and non-human primates shares properties with VTE, with returns to previously sampled objects associated with better future memory (Voss et al., 2011), and better performance during difficult perceptual discriminations (Voss & Cohen, 2017). Less evidence exists for VTE (based on head and body movements) in humans during decision-making. However, one recent study (Santos-Pata & Verschure, Paul F. M. J., 2018) found that VTE-like head movements were enhanced early in learning on a virtual maze, and were elevated specifically at an early (high-cost) decision point, similar to VTE behaviors in rats trained on a multiple T-maze (van der Meer, M. A., Johnson, Schmitzer-Torbert, & Redish, 2010). As VTE is an important behavioral index of deliberation in rodents (as reviewed in Redish, 2016), the existence of VTE in human navigation, and the relationship of VTE in humans to deliberation (in tasks such as the Web-Surf) is an important issue for translational research on decision-making.

In order to compare behavioral measures of decision-making during navigation in humans to that of rodents, we developed two virtual navigation versions of the Restaurant Row task for humans (here referred to as the *Movie Row* and *Candy Row*). The tasks were presented on standard desktop and laptop computers. In the task, participants navigated on a square track, similar to that used in the rodent Restaurant Row, using the keyboard arrow keys. At each reward site, participants were offered short video clips, taken from the Web-Surf (for the Movie Row), or candy/snacks (for the Candy Row), which were available after a variable delay. We predicted that behavior on the Movie Row and Candy Row tasks would replicate the published behavioral findings from the Web-Surf and Restaurant Row. We further expected that measures derived from navigation, including measures analogous to behaviors seen in rodents during VTE, would be associated with deliberation during difficult offers. To quantify VTE, we focused on behaviour observed within an offer zone, where participants made their decisions to accept or reject an offer for a specific reward. The primary outcome measures were decision latency (time from when the offer is presented until participants commit to accept or reject the offer by leaving the offer zone), the total amount of rotation (analogous to the rodent measure of VTE, the integrated absolute angular change in the orientation of motion of the head, Papale, Stott, Powell, Regier, & Redish, 2012), reversals in rotation direction, total distance travelled, and cumulative time spent paused in the offer zone. We predicted that during difficult decisions, when the delay offered was close to one’s stay versus skip threshold, the latency to make a decision would be elevated, and participants would be more likely to make one or more corrections before committing to their final choice.

We also conducted a set of exploratory analyses, to determine the relationship between measures of delay discounting for money (the Monetary Choice Questionnaire, MCQ, Kirby, Petry, & Bickel, 1999) and food (the Food Choice Questionnaire, FCQ, Hendrickson, Rasmussen, & Lawyer, 2015) to behavioral measures derived from the Movie and Candy Row tasks. We predicted that discounting rates would be related across the two surveys, and that willingness to wait for rewards would be related across the two navigation tasks. Based on previous work with a version of the Web-Surf involving risk (Abram, Redish, & MacDonald III, 2019), we predicted that discounting rates would not be related to performance on the foraging tasks.

From a subset of participants, information about smoking status and BMI were available from screening surveys. A final set of analyses were conducted to replicate previous reports that smoking and obesity are associated with stronger delay-discounting and to explore the relationship between smoking status and obesity to behavior on the Movie and Candy Row tasks.

## Method

### Participants

Undergraduates (male, n = 144), workers from Amazon’s Mechanical Turk service (mTurk, n = 147, 80 female/66 male/1 non-binary), and members of the local community (n = 34, 19 female/15 male) participated in the study. Undergraduates received course credit, extra credit or coupons for a campus coffee shop ($10) for participation, while mTurk workers were paid between $8-$8.50 USD for completing one version of the Movie Row task, and those recruited for the second sample also received $0.15 USD for completing a demographics survey, and $2.50 USD for completing a screening questionnaire. Participation by mTurk workers was limited to individuals in the United States, and those who had an approval rate for previous assignments of at least 90%. Community members were recruited by direct mail, online advertisement (Facebook, Craigslist) and public online classified ads posted on the Wabash College website, and received a $6 gift card for completing an initial survey battery, and $25 for travelling to campus and completing both the Movie and Candy Row tasks.

## Materials

### Movie Row task

The computer task was implemented using the Unity 3D game engine (http://unity3d.com), embedded in a web page. Data collected by the task were saved in a MySQL database, submitted using custom PHP scripts. Four versions of the task were tested as the task was refined across data collection in the context of undergraduate research projects (testing the impact of glucose consumption, dieting/obesity, and smoking on decision-making). Details on the samples recruited for these studies and links to the current version of the Movie Row are provided in the Supplementary Online Material.

#### Version 1

Participants navigated in a virtual environment, on a rounded square track. Four movie screens were positioned on the corners of the track, each of which played a different type of movie (kittens, bike accidents, dancing, and scenes of landscapes: movies were taken from Abram et al., 2016). Examples of each movie type are shown in Figure 1C, top. The movie type assigned to a given screen was randomly determined for each participant. Participants entered an offer zone as they approached the screen (visible as a line on the track, though not explicitly described to the participant), where they were presented with the offer (the type of video, and the delay). To accept the offer, they were required to step onto a circular platform, and look at the movie screen. If they looked away, the delay would pause, and would resume if they looked back at the screen. Participants could also skip movies entirely or quit during the delay (choice options were thus comparable to those in Sweis et al., 2018).

**Figure 1.**
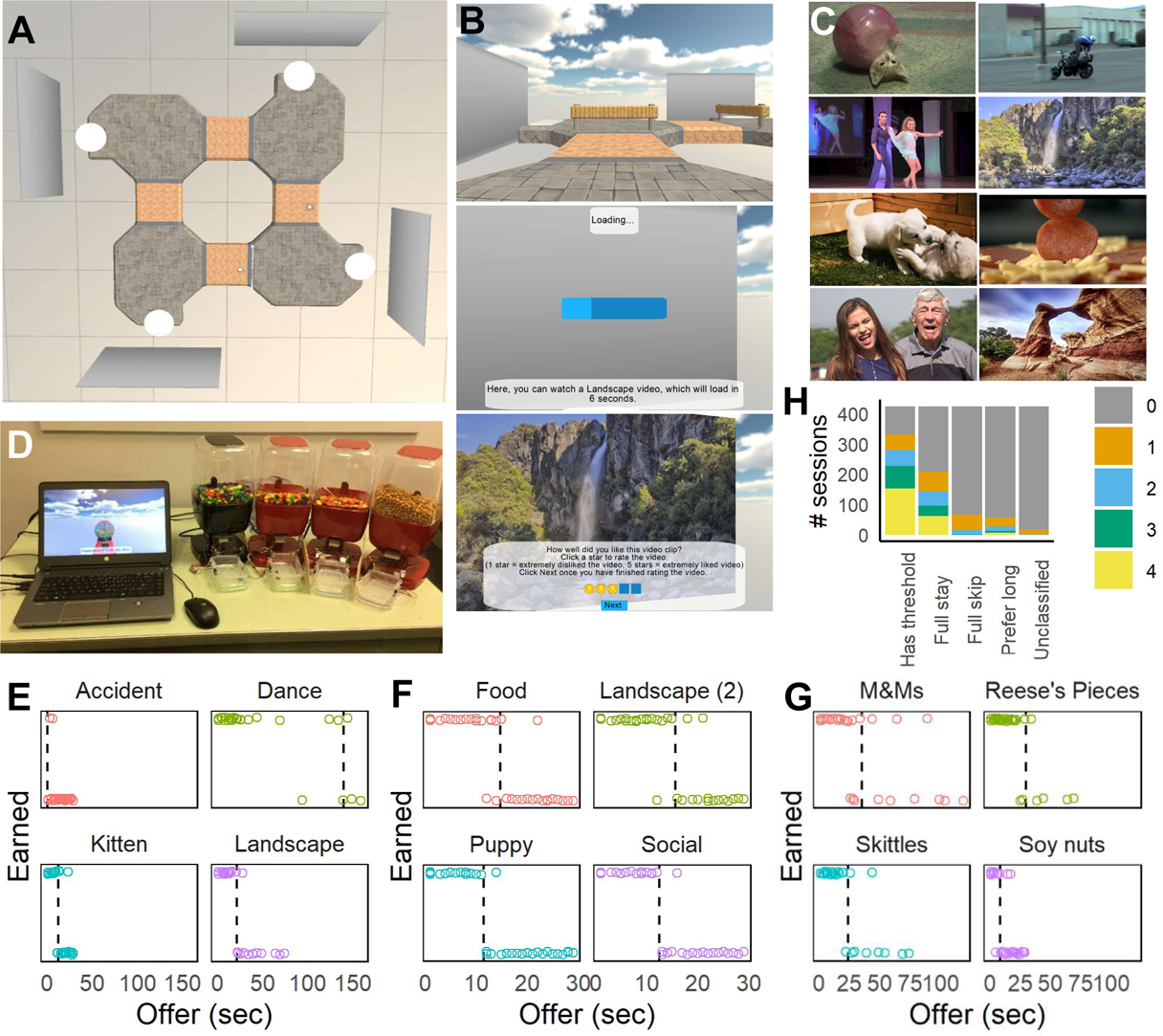
Virtual navigation tasks and choice behavior. A: Overhead view of the Movie Row track used after Version 1. Four movie screens were used, each displaying a different type of video clip. B: First-person screenshots taken during the task, showing navigation between offer zones (top), loading bar as the participant waits for an offer (middle), and showing the post-video rating after a video clip is played (bottom). C: Examples of movies used in the Movie Row task from the Web-Surf (top: kittens, accidents, dancing, and landscapes) and a second set (bottom: puppies, food, social, landscapes). D: Setup for the Candy Row task and food dispensers. Movie screens in the task were replaced with virtual candy (gumball) dispensers. E-G: Examples of choice behavior for one participant (DDR035) tested on the Movie Row task with the original (E) and new (F) videos, and on the Candy Row task (G). Sessions from E & G were done in the lab, and the session from F was done by the participant online. H: Distribution of the types of responding (number of delay thresholds, full-stay or full-skip behavior and preferences for long delays) across the sample.

The task continued until a participant had received an offer for each delay for each movie type, or until 40 minutes in the main task had elapsed. Most finished much faster (about 30 minutes), unless they watched the majority of the movies. Each video clip lasted 4 seconds, and were accompanied by audio. Headphones were provided for participants tested in-person in groups in a computer lab. Movies were randomly selected from the set of those available for each type, and used without replacement. Delays (3 to 29 seconds) were randomly selected for each type, also without replacement. The task began with a practice phase, where participants were given extra instructions to guide them through the task, and were allowed to choose and skip movies until they had watched two videos of the same type (requiring at least one full circuit of the track). After completing the task, participants completed a post-task survey asking them to rank the videos from 1 = favorite to 4 = least favorite.

#### Version 2

In the second version tested, the practice phase was revised so that participants were required to skip and watch one video of each type (the order randomized for each movie screen). If participants accepted an offer that was within 5 seconds of the maximum delay, the maximum delay for that movie type was increased by 5 seconds (in order to better estimate delay thresholds when participants were willing to wait longer than 29 seconds for some video types). The track was revised as well (see Figure 1A) to refine the offer zone, requiring participants to turn approximately 90-degrees to their left to accept a video, and to turn 90-degrees to their right to skip a video and continue to the next video type. Participants also completed a post-task survey including the rankings of each reward type, asked about their ability to hear the audio, and asked participants to describe how they selected which videos to watch and skip.

#### Version 3

To assess reaction time, a gate was placed at the entrance to each offer zone. As participants arrived, their avatar was frozen until the gate lowered (over a window of one second) and the offer was presented. Reaction time was measured from the time the avatar was unfrozen until the participant started moving.

#### Version 4

While participants typically accepted offers with short delays, and skipped offers with long delays, occasionally participants demonstrated the opposite pattern and were more willing to wait for long delays. This behavior was more common in online samples, and occasionally was associated with reports by participants that they preferred long delays so that they could switch to another task during the delay. In an attempt to decrease this behavior, a fixation task was implemented during the delay, where two-digit numbers (between 10-99) were presented at random intervals during the delay. Once the delay was complete, participants were presented with four different two-digit numbers as options on the screen, and required to select the most recently presented number. After correct choices, the movie would play immediately. After incorrect responses, the message “Incorrect” was presented, and a 3 second time-out was implemented before the movie was started.

To facilitate the process of accepting an offer, the platform location was also expanded (to encompass the majority of the area outside of the offer zone), and a wider range of viewing angles (to the screen) were allowed to start the delay timer. With these changes, the delays would often start as soon as participants left the offer zone. However, this allowed participants to complete the delay from positions where the movie screen was only partially visible, or visible at an angle. So, videos were displayed immediately in front of the participant on a screen that was visible only when rewards were presented (rather than on the virtual screen used in earlier versions).

In the test phase, participants received offers with 1-second delays for the first 3 visits to each reward site, to provide participants with more experience with each reward before longer delays were implemented, and to better determine if participants who skipped the majority of offers for a video type would be willing to wait for even very short delays. For version 4, some participants (n = 44) were tested with an alternate set of movies, including scenes of food, puppies, social interactions and landscapes (Figure 1C, bottom). These videos were provided by the laboratory of Aoife O’Donovan, and were selected from YouTube videos that were high quality (> 1080HD), relevant the category, and perceived as rewarding by the researchers. Any text visible in the videos was removed by cropping.

### Candy Row task

The Movie Row task was adapted to deliver physical rewards (candy and snacks) using four commercial motion activated candy dispensers (Motion Activated Snack Dispenser, discontinued, Sharper Image) which delivered one of four rewards: M&Ms, Skittles, Reese’s Pieces or soy nuts/pepitas (see Figure 1D). The program was presented on a laptop computer, in full screen mode, rather than in a web page. Participants were tested individually either alone in a small room or in a classroom under the supervision of a research assistant. While the three candy rewards remained constant throughout all of the testing, the fourth option varied across participants, due to compatibility issues with the dispensers. Goldfish crackers, peanuts and Cheerios were also used with some participants, and for this reason, the fourth reward is described here as the “other” option. At each offer zone, the movie screens were replaced with a virtual gumball dispenser. Instructions were modified accordingly, describing each zone as a candy shop. Delays were described as the time required to prepare the reward for delivery, rather than the loading time (as in the Movie Row task).

Dispensers were controlled using an Arduino running custom software to interface with the virtual navigation task. Each dispenser was fitted with an infrared detector to verify reward delivery, and rewards were delivered into small plastic magazines, which were also fitted with infrared detectors (to assess reward retrieval by participants).

Participants initially were allowed to sample each of the four rewards (presented outside of the task), and asked to rate their enjoyment of the reward on a scale of 0 to 6 (0 = Not at all, 2 = Slightly, 4 = Quite a bit, 6 = Extremely), and to rank the four rewards from favorite to least favorite. After the task, participants also completed a post-task ranking of the rewards and asked participants to describe how they selected which food offers to accept and skip, similar to the Movie Row task. Rewards that participants earned, but which were not consumed, were weighed and given to the participant after completion of the task.

### Delay-discounting surveys

#### Monetary Choice Questionnaire (MCQ)

A 21-item version of the Monetary Choice Questionnaire (Kirby et al., 1999) was administered during a set of screening and initial surveys, using the reward magnitudes and delays used by Wang, Reed, Baugh and Fercho (2018). For each item involving hypothetical monetary rewards available immediately or after a delay, participants selected either to accept a small immediate offer, or to wait the specified time for a larger reward.

#### Food Choice Questionnaire (FCQ)

The 27-item version of the Food Choice Questionnaire (Hendrickson et al., 2015) was administered for a subset of participants before the MCQ. For each item, participants chose between hypothetical food rewards of different magnitudes (specified as a number of bites of one’s favorite food). In the original description of the survey, participants were presented with a physical cube to represent the size of one bite (5/8 of an inch, or 1.59 cm). For online administration, the survey was adapted to omit this physical cue, and participants were instructed to “*Please imagine that each bite of food is equal to the size of a cube that is half of an inch tall (about 1.25 centimeters tall).*”

### Procedure

Participants were tested in-person at Wabash College (for some Movie Row sessions and all Candy row session) or online (some Movie Row sessions). Participants tested on the Movie Row in the laboratory completed the task in a computer lab on PCs divided by partitions. Headphones were provided for the Movie Row, so that participants could hear the audio presented with each video.

For some participants, information on body mass index (BMI), smoking status, and survey measures of delay discounting for money (using the values and delays from Wang et al., 2018) and food, were available from surveys completed before the Movie Row or Candy Row tasks. BMI values were calculated from self-reported weight and height. Using the Center for Disease Control criteria (www.cdc.gov/obesity/adult/defining.html), values were classified as underweight (BMI < 18.5), healthy (18.5 ≤ BMI < 25), overweight (25 ≤ BMI < 30) or obese (BMI ≥ 30), and analyses using BMI were conducted comparing underweight/healthy individuals (BMI < 25) to overweight/obese individuals (BMI ≥ 25).

### Analysis

#### Delay thresholds

The participants’ threshold for deciding to watch or skip videos of each type was estimated by fitting their decisions to stay (= 1) or skip/quit (= 0) each reward as a function of the delay offered on each trial. Decisions were padded with one skip (0) at 1 second less than the minimum delay offered and one stay (1) at 1 second longer than the maximum delay offered, to account for cases where participants accepted or rejected all of the offers. Fits were performed using a Heaviside function, using a leave-one-out approach in which a threshold for each trial was calculated using every other offer for the same reward type, to calculate the trial-specific threshold (Abram et al., 2019). Analyses of value used the difference between the offer on the current trial and the trial-specific delay threshold. The average across the trial-specific delay thresholds within a reward type was used as the participant’s overall delay threshold for each reward category. To account for cases in which participants preferred long delays, rather than short delays, the Heaviside function was also fit to the inverse of the choice behavior. The error (number of trials that deviated from the predicted outcome) was calculated for both fits. A participant was defined as having a normal delay threshold for a reward category if the average error associated with the Heaviside fit was lower than that of the inverse fit. A participant was defined as having a preference for long delays if the error associated with the Heaviside fit was higher than that of the inverse fit. In the special case where participants skipped/accepted all or nearly all of the offers, the error of both fits would be equal, and these special cases were classified as full-stay (if participants accepted at least 75% of all of the offers for the reward type) or full-skip (if the participant skipped at least 75% of all of the offers for the reward type). For all other cases, the reward type was considered to be unclassified.

#### Magazine entries

Breaks of the magazine infrared beam were used to assess retrieval of rewards in the Candy Row. Breaks lasting between 50 milliseconds and 10 seconds were scored as a retrieval, and the number of retrievals and duration of the entry were recorded for each trial in each magazine. As participants often did not consume all of the rewards earned, and the infrared sensor could become obstructed by the excess foods, analysis of magazine entries was restricted to the first 20 minutes of the test session.

#### Offer zone

After version 1 of the Movie Row task (which lacked a clearly defined offer zone), several measures were calculated for each trial on behavior observed from the time the offer was presented (as the participant entered the offer zone), until the participant crossed out of the offer zone (moving on to the next reward zone, or moving towards the platform). The measures examined were latency to leave the offer zone, distance travelled in the offer zone, total rotation, pausing, rotation reversals, and entry bias (position of the participant when entering the offer zone, relative to the hallway center). Behavior after re-entry to the offer zone on a single trial were not included in these measures. Offer zone behavior on trials with decision latencies longer than 15 seconds were removed. Behavioral measures (except rotation reversals) were normalized by first using a log^10^ transformation to remove a strong positive skew (toward long times/distances/rotations), then z-scoring (within session) to remove relationships to factors such as gender and age (see Supplementary Figure 1B).

#### Decision latency and distance travelled

Decision latencies were defined as the time spent in the offer zone (after the participant started moving in Versions 3-4, in milliseconds), and the integrated distance travelled until the participant left the offer zone (in arbitrary units).

#### Total rotation and rotation reversals

The amount of rotation in the offer zone was calculated as the sum of the difference in heading direction for all position samples from when the offer was presented until the participant left the offer zone. The number of reversals of rotation direction in the offer zone was also scored. Rotation measures on trials in which participants rotated more than 360 degrees in the offer zone were removed.

#### Pausing

Pausing in the offer zone was defined as the total amount of time (in milliseconds) participants stopped moving (no position changes or rotation).

#### Entry bias

After Version 1, entry bias was assessed as the location where participants entered the offer zone, relative to the middle of the pathway (defined as an entry bias of 0). This entry was scaled to the width of the hallway, and values ranged between −50% (towards the participant’s left and the reward location, 50% of the hallway width) to +50% (towards the participant’s right and the exit used to skip the reward).

#### Reaction time

In Versions 3-4, the time from when the offer was presented until the participant began moving was defined as the reaction time. Reaction times were log_10_ transformed and z-scored within-session. Data was not analyzed for trials where this measure exceeded 15 seconds.

#### Regret

Following the procedure of Steiner and Redish (2014), trials were classified as regret-inducing when participants were presented with an offer that was above their delay threshold, after skipping or quitting an offer at the previous zone which was below their delay threshold (thus receiving a low-quality offer after passing up a higher quality offer). These regret trials were compared to two control conditions, Control-1: when participant received an offer that was above their delay threshold, and had previously accepted an offer which was below the delay threshold for the previous zone, and Control-2: when participant received an offer that was above their delay threshold, and had previously skipped or quit an offer which was above the delay threshold for the previous zone.

#### Sunk-cost analysis

Following the procedure of Sweis and colleagues (Sweis et al., 2018), the effects of sunk-costs were estimated using relationship between the probability of completing a trial (once the loading bar was started) as a function of the amount of time remaining in the delay. Across participants, the percentage of trials in which participants waited the entire duration of the delay was calculated as a function of the initial delay (for offers between 1 and 29 seconds). To assess the impact of sunk-costs, the likelihood of completing a given delay was compared to trials in which participants had invested some amount of time (making an investment by waiting a minimum amount of time). For example, to estimate the effects of sunk costs after participants had invested 5 seconds, all trials in which participants started the delay and waited at least 5 seconds were selected. Then, the percentage of trials in which participants completed the remainder of the delay was calculated, as a function of the amount of time remaining, over delays from 1 to 23 seconds (corresponding to initial offers of 6 to 29 seconds, see Figure 5A).

## Results

### Relationship to delay discounting measures

Two survey measures of delay discounting were administered before the Movie/Candy Row tasks, the Monetary Choice Questionnaire and the Food Choice Questionnaire. Each involved hypothetical monetary or food rewards of three magnitudes (small, medium, large). We expected that delay discounting rates would decrease as a function of reward magnitude, and would also be higher for smokers and those who report high BMIs (> 25, falling in the overweight or obese range, Jarmolowicz et al., 2014). Responses for both the monetary and food versions of the survey were orderly, judged by the percentage of responses for each participant which were consistent with their estimated discounting rate (k). The mean percentage of responses consistent with estimated k values for the MCQ (for small (98%), medium (99%) and large (99%) reward magnitudes) were slightly higher than for the FCQ (for small (93%), medium (94%) and large (94%) reward magnitudes), but were both consistent with those reported by Hendrickson, Rasmussen and Lawyer (2015).

An ANOVA was used to analyze discounting rates (log_10_ k) for the MCQ with Magnitude (small, medium, large) as a within-subjects factor, and Gender (male, female) as a between-subjects factor. Discounting rates decreased as the reward size increased (F(2, 1526) = 1,726, p < 0.001, η^2^_p_ = 0.69), while neither the main effect of Gender (F(1, 763) < 1, n.s., η^2^_p_ < 0.001) nor the Gender × Magnitude interaction were significant (F(2, 1526) = 1.2, p = 0.29, η^2^_p_ = 0.002). For the subset of participants who completed the FCQ, there was also a main effect of Magnitude (F(2, 576) = 3.5, p = 0.03, η^2^_p_ = 0.012), which was qualified by a significant Gender × Magnitude interaction (F(2, 576) = 9.9, p < 0.001, η^2^_p_ = 0.033), while the main effect of Gender was not significant (F(1, 288) = 3.0, p = 0.86, η^2^_p_ = 0.01). Post-hoc tests (Bonferroni corrected, α = 0.05) revealed that for the FCQ, discount rates for males for small magnitude rewards were significantly higher than for medium and large rewards, but for females there were no differences across the three reward magnitudes. While responses to the FCQ showed less sensitivity to reward magnitude in general, overall discounting rates (log_10_ k) were positively correlated in the subset of participants who completed both the MCQ and FCQ (r(288) = 0.22, p < 0.001, see Table 1 and Figure 2D, left), indicating that participants who were more willing to wait for hypothetical monetary rewards were also somewhat more willing to wait for hypothetical food rewards.

**Figure 2.**
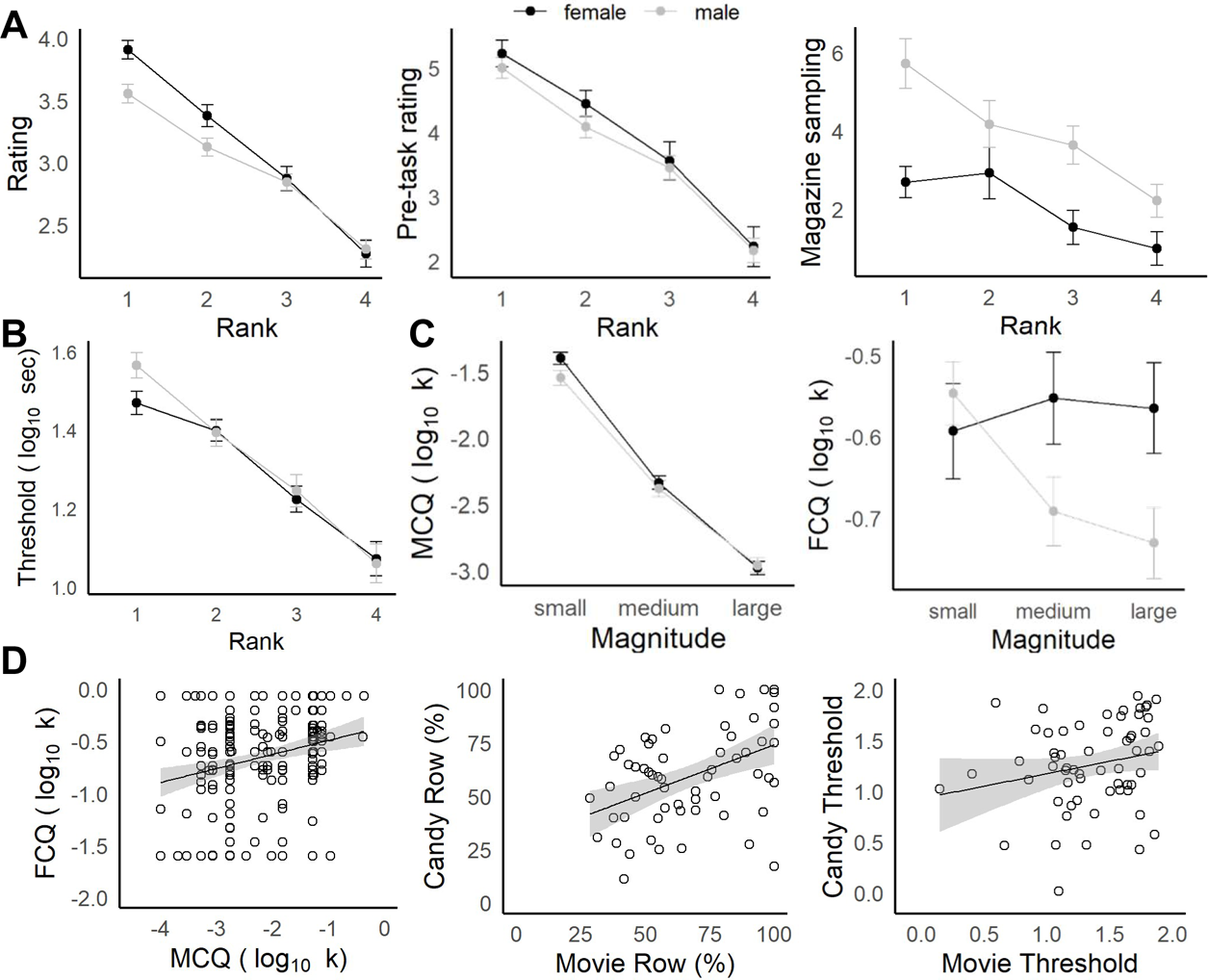
Characterizing navigation tasks and comparison to delay-discounting. A: Stated versus revealed preferences. Average star ratings (left) in the Movie Row, and pre-task enjoyment ratings (middle) in the Candy Row decreased as a function of the post-experiment ranking. Participants were more likely to retrieve their favorite food rewards on the Candy Row (right). B: Delay thresholds decreased significantly across ranks (favorite to least favorite). C: Delay-discounting (k) for hypothetical money rewards (left) did not differ by gender, and decreased as a function of reward magnitude. For hypothetical food rewards (right), males showed a similar sensitivity to magnitude, while females did not. D: Comparison of navigation tasks and delay-discounting measures. Overall delay-discounting for hypothetical money and food rewards was correlated (left) in the survey measures, as was the number of actual food and movie rewards earned (middle) and a similar trend was seen for delay thresholds (right).

**Table 1.**
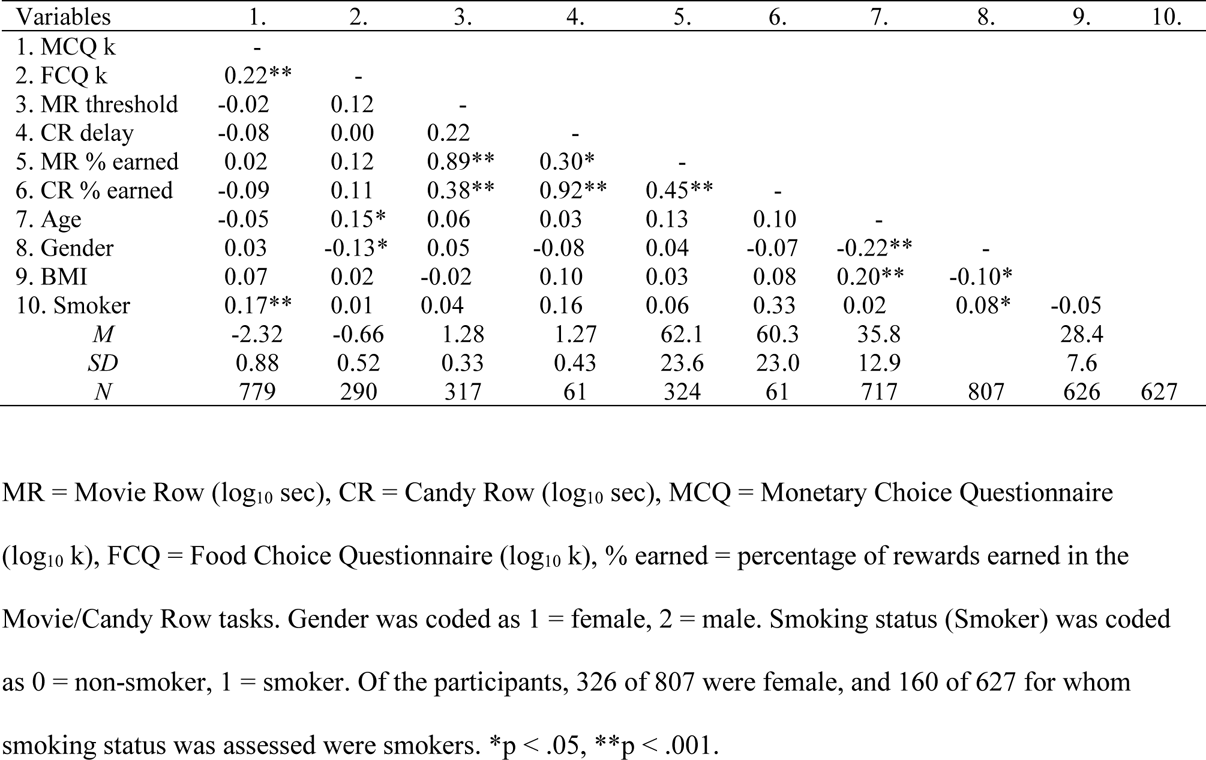
Correlation between discounting rates (log_10_ k) and row task performance.

In separate ANOVAs, overall discounting rates (log_10_ k) for the MCQ and FCQ were analyzed, with Gender, Smoker (non-smoker, smoker) and BMI (<25: underweight/healthy, >25: overweight/obese) as between-subjects factors. For the MCQ, smokers had significantly higher discounting rates compared to non-smokers (F(1, 513) = 17.8, p < 0.001, η^2^_p_ = 0.034, see Figure 3A), and overweight/obese individuals tended to have higher discounting rates compared to underweight/healthy individuals (F(1, 513) = 3.6, p = 0.059, η^2^_p_ = 0.007, see Figure 3B). No other main effects or interactions were significant (p > 0.24, η^2^_p_ < 0.003). For the FCQ, no main effects or interactions with smoking status or BMI group were significant (p > 0.12, η^2^_p_ < 0.012).

**Figure 3.**
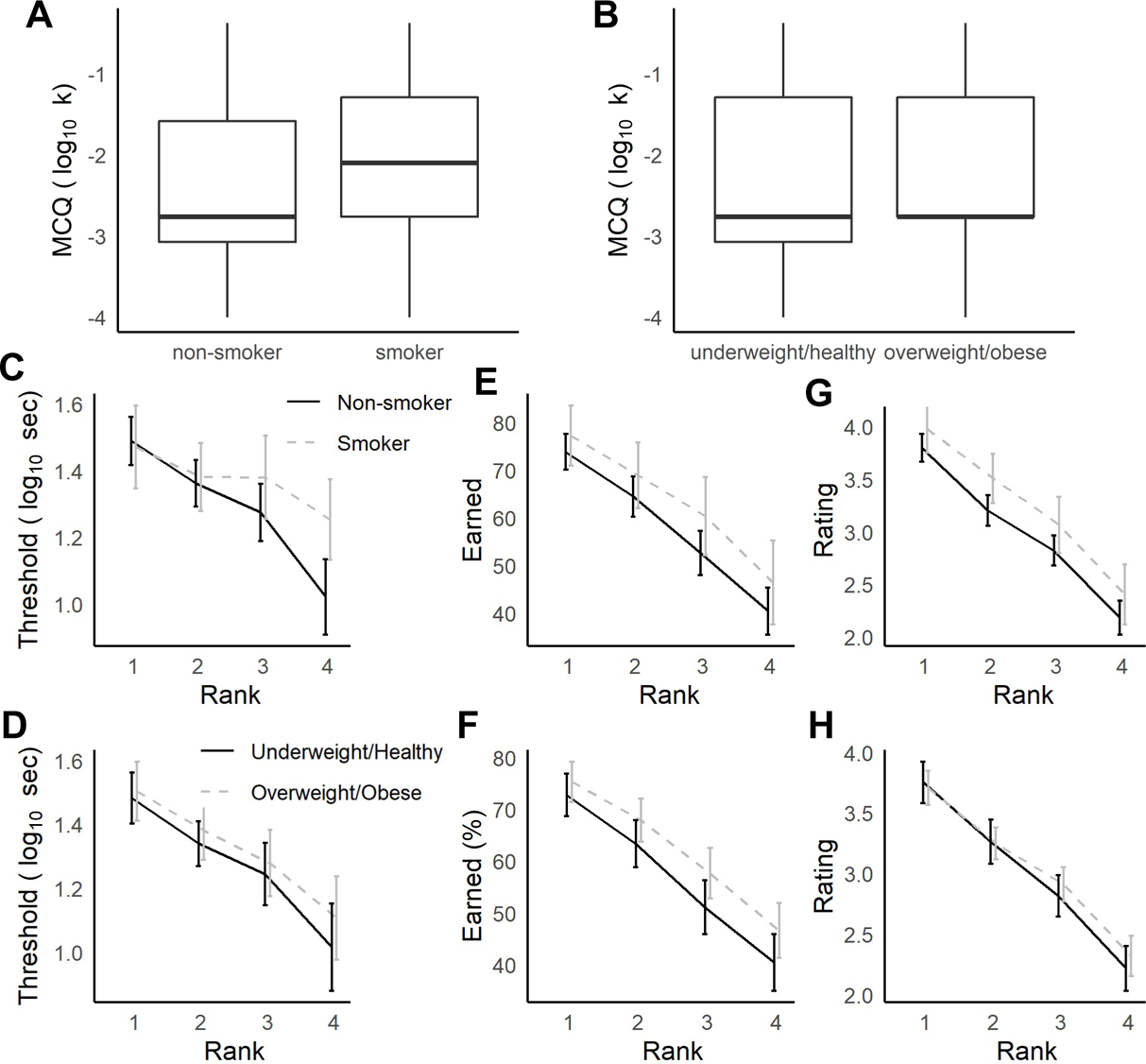
Relationships between delay-discounting and foraging measures to smoking status and obesity. A-B: Smokers more strongly discounted hypothetical money offers in the MCQ (A), and overweight/obese individuals tended to do so as well (non-significant). C-H: During foraging, delay thresholds were not strongly related to smoking (C) or BMI (D). Smokers (E) and overweight/obese individuals (G) tended to earn more rewards (non-significant), but in smokers this trend could be explained by higher subjective ratings of the rewards on the Movie Row task (G), while BMI group was not related to movie ratings (H).

### Row tasks

Across the participants who completed the Movie and/or Candy Row tasks (and received at least 40 offers in the test phase, n = 326), many participants demonstrated the expected pattern of accepting offers with short delays and rejecting offers with long delays. And, many participants demonstrated a preference across the four reward types, such that they were more willing to wait for some types of movies or foods (e.g. Figure 1E-G). In some cases, participants accepted the majority or very few of the offers of a given reward type (see example in Figure 1E for Accident videos). And, some participants showed the opposite of the expected pattern (preferring rewards offered at long delays). To examine the proportion of participants showing each response pattern, each participant’s responses to each reward type were classified as having a delay threshold, preferring long delays, full-stay, full-skip, or unclassified. The number of reward types (between 0 to 4) falling into each classification were counted for each participant, and the distributions for each sample are shown in Figure 1H.

In first sample tested (using version 1, and tested in a laboratory), the most common response pattern was to demonstrate a delay threshold for at least one video type. However, a sizable fraction of participants also showed full-stay behavior for 3-4 of the movie types. Informal feedback from participants suggested that some were unaware that they could decide which videos to watch and skip. To address this potential concern, in version 2 the instructions and practice phase were refined as described in the Methods. However, a fraction of participants tested online (with versions 2 and 3) had at least one movie type classified as full-stay or full-skip, and some also demonstrated a preference for long delays. Some of these differences are likely attributable to the use of online data collection, in which it was difficult to ensure that participants stayed on-task during the delay before the video loaded. In fact, some participants indicated (in the post-task survey) that they used the time during the delay to do other tasks (and so long delays may have been more valuable to the participant than short delays). In samples tested in person in a laboratory with the later versions (3 and 4), the task performed well (with 83% of participants showing at least one delay threshold).

### Stated preferences across video types

Participants provided self-report measures of their stated reward preferences (the ratings given immediately after watching each video or the pre-task ratings of each food, and the post-task rankings of the video/food types from most to least favorite). Additionally, in the Candy Row task another measure of reward preference was the frequency of sampling the reward magazine (to retrieve earned rewards). Abram et al. (2016) reported a significant interaction between gender and stated preferences for the four movie types in the Web-Surf task, and similar results were obtained for the Movie Row (see Supplementary Figure 1A). For the videos from the original Web-Surf task, using a two-factor ANOVA with Gender (female, male) as a between-subjects factor and Reward Type (Kitten, Accident, Landscape and Dance) as a within-subjects factor, there was a main effect of Reward Type (F(3, 855) = 83.9, p < 0.001, η^2^_p_ = 0.23), and no main effect of Gender (F(1, 285) = 1.7, p = 0.19, η^2^_p_ = 0.006). As predicted, the interaction of Reward Type × Gender was significant (F(3, 855) = 133.0, p < 0.001, η^2^_p_ = 0.10), with females giving higher ratings to Kitten videos than did males, and males giving higher ratings to Landscape videos than did females. Average post-video ratings did not differ significantly by gender for Accident and Dance videos.

In a separate ANOVA for the second set of movies (Supplementary Figure 1B), there was a main effect of Reward Type (Puppies, Social, Food and Landscape, F(3, 105) = 15.6, p < 0.001, η^2^_p_ = 0.31), and no main effects of Gender (F(1, 35) < 1, n.s., η^2^_p_ = 0.02) nor an interaction of Reward Type × Gender (F(3, 105) < 1, n.s., η^2^_p_ = 0.008). Similar results were found for the Candy Row task (Supplementary Figure 1C-D) for pre-task enjoyment ratings given to the food options, where there was a main effect of Reward Type (M & Ms, Reese’s Pieces, Skittles and Other, F(3, 192) = 6.2, p < 0.001, η^2^_p_ = 0.09), and no main effects of Gender (F(1, 64) < 1, n.s., η^2^_p_ = 0.008) nor an interaction of Reward Type × Gender (F(3, 192) < 1, n.s., η^2^_p_ = 0.01). For magazine entries, there was also a main effect of Reward Type (F(3, 171) = 3.6, p = 0.015, η^2^_p_ = 0.06), while the Type × Gender interaction was not significant (F(3, 171) < 1, n.s., η^2^_p_ = 0.008). Males did make more magazine entries on average than females (F(1, 57) = 4.8, p = 0.032, η^2^_p_ = 0.08) across the four reward types.

### Relationship between stated and revealed preferences

Revealed preferences were assessed by the delay threshold for each reward type, and by the proportion of offers that were accepted for each reward type. To examine the relationship between stated and revealed preferences, the reward types were sorted (recoded) based on the post-task rankings (from most to least favorite) for each participant. Separate ANOVAs were conducted to examine the relationship between Rank (1-4, favorite to least favorite) and Gender on the average post-video rating (in the Movie Row task), enjoyment ratings and magazine sampling (in the Candy Row task), the delay thresholds, and the proportion of offers accepted.

As expected, there was a strong correspondence between the measures of stated preference (see Figure 2A). Using ANOVAs with Rank (1-4) as a within-subjects factor and Gender as a between-subjects factor, there was a significant main effect of Rank for the post-video ratings in the Movie Row for both the first (F(3, 891) = 310, p < 0.001, η^2^_p_ = 0.51), and second (F(3, 891) = 310, p < 0.001, η^2^_p_ = 0.51), sets of videos, and on the Candy Row task for pre-task enjoyment ratings (F(3, 192) = 80, p < 0.001, η^2^_p_ = 0.56). In the Candy Row, there was also a main effect of Rank for magazine sampling (F(3, 171) = 10.6, p < 0.001, η^2^_p_ = 0.16). Across the four ranks, post-video ratings, pre-task enjoyment ratings, and the likelihood of retrieving a food reward earned decreased strongly.

Additionally, there was also a Rank × Gender interaction for post-video ratings for the original Web-Surf videos (F(3, 891) = 7.2, p < 0.001, η^2^_p_ = 0.024) and a significant main effect of Gender for magazine sampling in the Candy Row task (F(1, 57) = 3.2, p =0.023, η^2^_p_ = 0.078), while other main effects were not significant (all Fs < 1.7, ps > 0.22, η^2^_p_ < 0.026). For the original Web-Surf videos, females gave significantly higher ratings to their top-ranked videos compared to males, while on the Candy Row task, males were more likely to sample the magazine (and presumably retrieve rewards).

The proportion of rewards earned and delay thresholds were also strongly related to post-experiment reward rankings: participants were more willing to wait for and earned significantly more of their top-ranked rewards, compared to their least favorite rewards (Figure 2B). In two separate two-factor ANOVAs (selecting the first session completed by each participant across the two tasks), there was a main effect of Rank (Proportion watched: F(3, 945) = 162, p < 0.001, η^2^_p_ = 0.34, Delay thresholds: F(3, 561) = 21.0, p < 0.001, η^2^_p_ = 0.10). For both measures, there were no significant main effects of Gender (Proportion watched: F(1, 315) = 1.1, p = 0.29, η^2^_p_ = 0.004, Delay thresholds: F(1, 187) = 1.6, p = 0.21, η^2^_p_ = 0.008) nor a Rank × Gender interaction (Proportion watched: F(3, 945) = 1.2, p = 0.30, η^2^_p_ = 0.004, Delay thresholds: F(3, 561) < 1, n.s., η^2^_p_ = 0.001). This pattern of results was not changed if the analysis was restricted to the Candy Row, or to one of the video sets used in the Movie Row task.

These results indicate that while females gave higher star ratings than males to their top ranked video type (stated preference) for the original Web-Surf videos, and males were more likely to reach into the food magazines in the Candy Row, there were no significant gender differences in revealed preferences (delay thresholds and the proportion of rewards earned).

### Relationship to delay discounting

Overall, the percentage of rewards earned were positively correlated in the subset of participants who completed both the Movie and Candy Row tasks, and a similar trend was observed for delay thresholds, averaged across the log_10_ transformed delay thresholds calculated within each reward type (percentage of rewards earned: r(58) = 0.447, p < 0.001, delay thresholds: r(58) = 0.221, p = 0.089, Figure 2D, middle and right), similar to results found for the MCQ and FCQ (see Table 1 and Figure 2D). However, delay thresholds and the percentage of rewards earned on the Movie and Candy Row tasks were not significantly related to delay-discounting rates derived from the MCQ and FCQ surveys (all ps > 0.11). These results indicate that willingness to wait for hypothetical monetary and food rewards in the survey measures was not predictive of actual willingness to wait for video and food rewards in the experiential foraging tasks.

### Relationship to smoking and BMI

While the overall discounting rates for the MCQ were elevated for smokers and a similar trend was observed for overweight/obese individuals, there were no strong differences observed in delay thresholds or proportion of rewards earned on the Row tasks among participants for whom smoking status and BMI were available (Supplementary Table 2). And, any trends observed in these samples were in the opposite direction, where smokers were more willing to wait for rewards on the Movie/Candy Row tasks (see Figure 3C-F). In separate three-way ANOVAs, used to analyze log_10_ transformed delay thresholds and the percentage of rewards earned, with post-task Rank (1-4) as a within-subjects factor, Gender as a between subjects-factor and either Smoker or BMI group as a second between-subjects factor, no effects of Gender or interactions were observed (ps > 0.45, η^2^_p_ < 0.003), nor any main effects or interactions of BMI group (ps > 0.27, η^2^_p_ < 0.017). Though the main effects for smoking status were not significant, smokers tended to have somewhat higher delay thresholds (F(1, 138) = 3.2, p = 0.076, η^2^_p_ = 0.023) and to earn more rewards (F(1, 175) = 3.7, p = 0.057, η^2^_p_ = 0.021).

These results indicate that, in contrast to the results for the MCQ, neither smoking status nor BMI group were significantly related to willingness to wait for rewards. And, any trends in these data suggested the smokers were willing to wait longer for actual video or food rewards in a foraging task. This trend, even if validated in a larger sample, may be driven by differences between smokers and non-smokers in their subjective enjoyment of the video rewards.

Analyzing the post-video ratings made on the Movie Row task using an ANOVA with Smoker and Gender as between-subjects factor, and Rank as a within-subjects factor, smokers gave significantly higher ratings to the videos they watched than did non-smokers (F(1, 172) = 3.9, p = 0.049, η^2^_p_ = 0.022). There was also a main effect of Rank (p < 0.001), but in the subset of participants for whom a smoking status was available, there were no main effects or interactions with Gender (ps > 0.14, η^2^_p_ < 0.012). Thus, it is possible that if smokers are somewhat more likely to accept offers on the Movie Row task, this could be driven by differences in enjoyment of rewards in the task.

### Regret

Steiner and Redish (2014) reported several behavioral correlates of regret in rats tested on the Restaurant Row, where in regret-inducing situations (receiving a poor offer after skipping a good offer), rats were more likely to accept a poor offer, spent less time consuming the reward, and were likely look back towards the previous (skipped) food location compared to the two control situations.

Compared to Restaurant Row, participants who completed Movie Row were less (rather than more) likely accept a poor offer when compared to the control conditions (Wilcoxon’s P, Control-1: p < 0.001, Control-2: p < 0.001, see Figure 4A). Additionally, participants were less likely to pause (Wilcoxon’s P, Control-1: p < 0.001, Control-2: p = 0.002), and compared to Control-1 trials, participants left the offer zone more quickly (Figure 6E, Wilcoxon’s P, Control-1: p < 0.001, Control-2: p =0.34), rotated less (Figure 6E, Wilcoxon’s P, Control-1: p < 0.001, Control-2: p =0.59), less often reversed rotation direction (Wilcoxon’s P: Control-1: p < 0.001, Control-2: p = 0.32), and travelled a shorter distance while in the offer zone, (Wilcoxon’s P, Control-1: p < 0.001, Control-2: p = 0.32). No significant differences in reaction time were observed between conditions (Wilcoxon’s P, Control-1: p = 0.37, Control-2: p =0.71). These data suggest that humans, similarly to rats, behaviorally differentiate between the regret-inducing and control trials, although there appear to be species or task-related differences (e.g. rats being more likely to accept offers on regret-inducing trials, and humans being less likely to do so).

**Figure 4.**
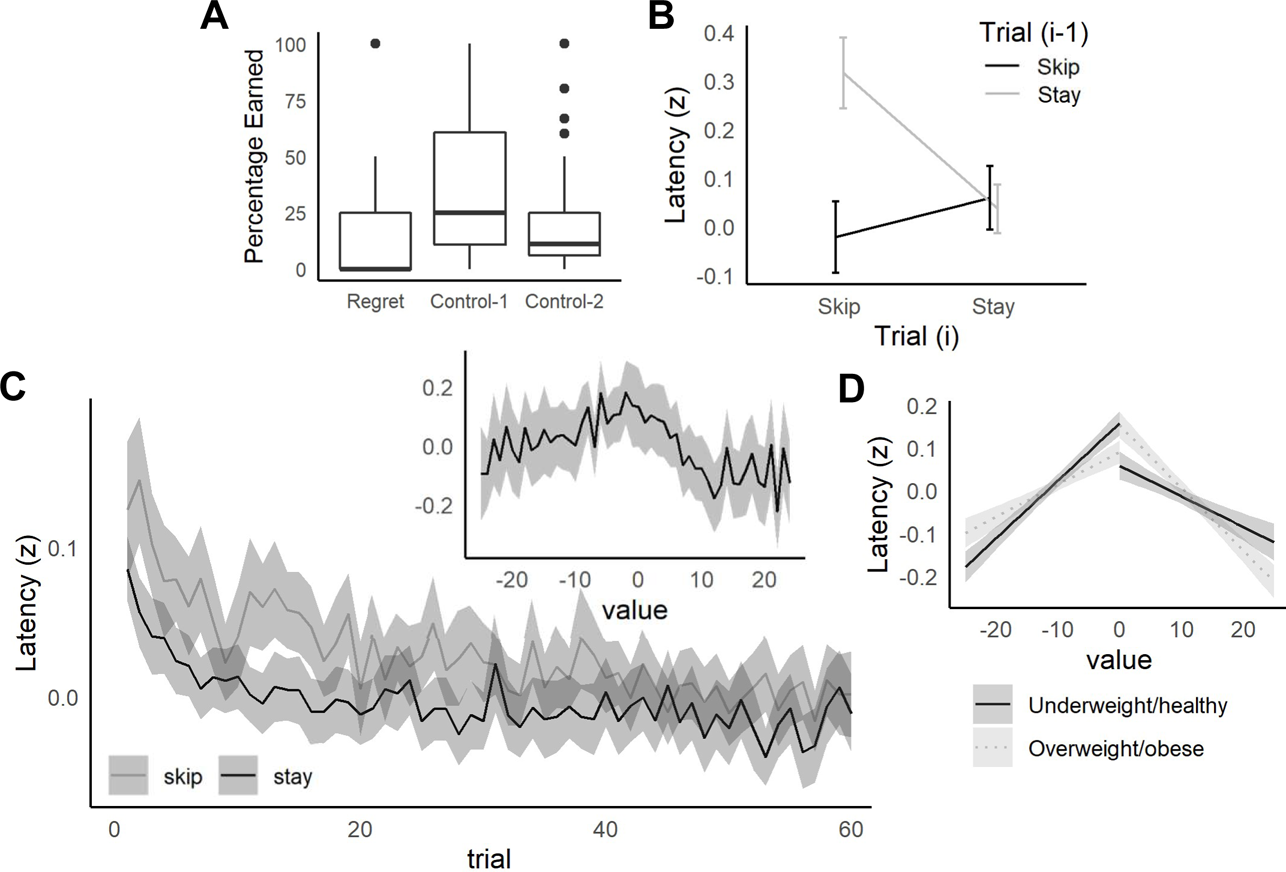
Replication of behavioral results from the Web-Surf. Regret. A: Participants were less likely to accept low-value offers (> threshold) in regret-inducing conditions (previously rejecting a low-value offer) compared to control conditions (control-1: previously accepted a high-value offer, control-2: previously skipped a low-value offer). B: Sequential choice behavior. Participants were slower to accept an offer (on trial i) after previously skipping an offer (on trial i-1), compared to other sequences (Skip/Skip, Stay/Stay, or Stay/Skip). C: Decision latency relative to trial number and value (threshold – delay). The time taken to make a choice (log-transformed and z-scored) was higher overall for skip decisions, but decreased across trials for both skip and stay decisions. Inset: Decision latency was also elevated for difficult decisions (value = 0, offers near threshold). D: Higher BMI individuals (> 25 – overweight/obese) were more sensitive to value for good offers (> threshold) compared to bad offers (< threshold). Lower BMI individuals (< 25 – underweight/healthy) tended to show a pattern in the opposite direction. Lines and shaded area represent estimated means and 95% confidence intervals based on the regression model. A-C: For each plot, data were averaged within participant, then means and confidence intervals were calculated across participants. Lines in C indicate means, and errorbars/shaded areas (B-C) indicate 95% confidence intervals.

### Sunk-costs

To determine if participants demonstrated a sunk-cost effect on the Movie and Candy Row, we examined how the probability of completing the entire delay for an offer was related to the amount of time already invested (as in Sweis et al., 2018). After starting the delay, participants on the Movie Row and Candy Row tasks were very unlikely quit compared to published data from the Restaurant Row (but were comparable to published data from the Web-Surf Task). Across the samples tested after version 1 of the Movie Row (which lacked a well-defined waiting zone), participants quit before the delay was completed on 0.7% of all trials in which participants stepped onto the platform (0.4% of all trials), which is lower than rates for both the Web-Surf and Restaurant Row. After reaching the platform, participants were highly likely to initiate the loading bar (in versions 2 and 3 of the Movie Row, which had a stricter requirement for starting the loading bar, participants started the loading bar on 99.6% of trials in which they stepped to the platform). Also, only 62 of the 296 participants (21%) tested after version 1 ever quit a trial after stepping onto the platform during their first Movie or Candy row task session. Among participants who quit on at least one trial after initiating the delay, participants quit on average 3.3% (SD = 2.3%, range = 1.5% to 14.3%) of trials in which they started the loading bar. The low quit rate on the Movie Row and Candy Row tasks may be related to the high response requirement to start the delay in versions 2-3 (participants must move to and stand on a small platform and look directly at the screen). However, quit rates remained low (M = 0.5%, SD = 1.1%) in version 4 and in the Candy Row, where the platform area was expanded and the requirement to initiate the delay was relaxed. It is also possible that low quit rates indicate a lack of awareness on the part of the participants that quitting during a delay was allowed (in spite of the instructions provided).

While quit rates were low on the Movie and Candy Row tasks, we did observe evidence that the decision to quit was sensitive to sunk-costs. As shown in Figure 5A, the likelihood of completing the delay decreased as a function of the initial delay (assessed across all participants, 95% CI for the slope = −0.00078, −0.00025). However, as participants have invested time (waiting during the delay), this relationship to delay weakens as a function of the amount of time invested (after completing 5 seconds, 95% CI = −0.00048, 0.00003, after 10 seconds, CI = −0.00023, 0.00013). While quit rates were low, once participants had invested several seconds waiting during a delay, they were more likely to complete the remainder of the delay, consistent with previous reports from the Web-Surf and Restaurant Row.

**Figure 5.**
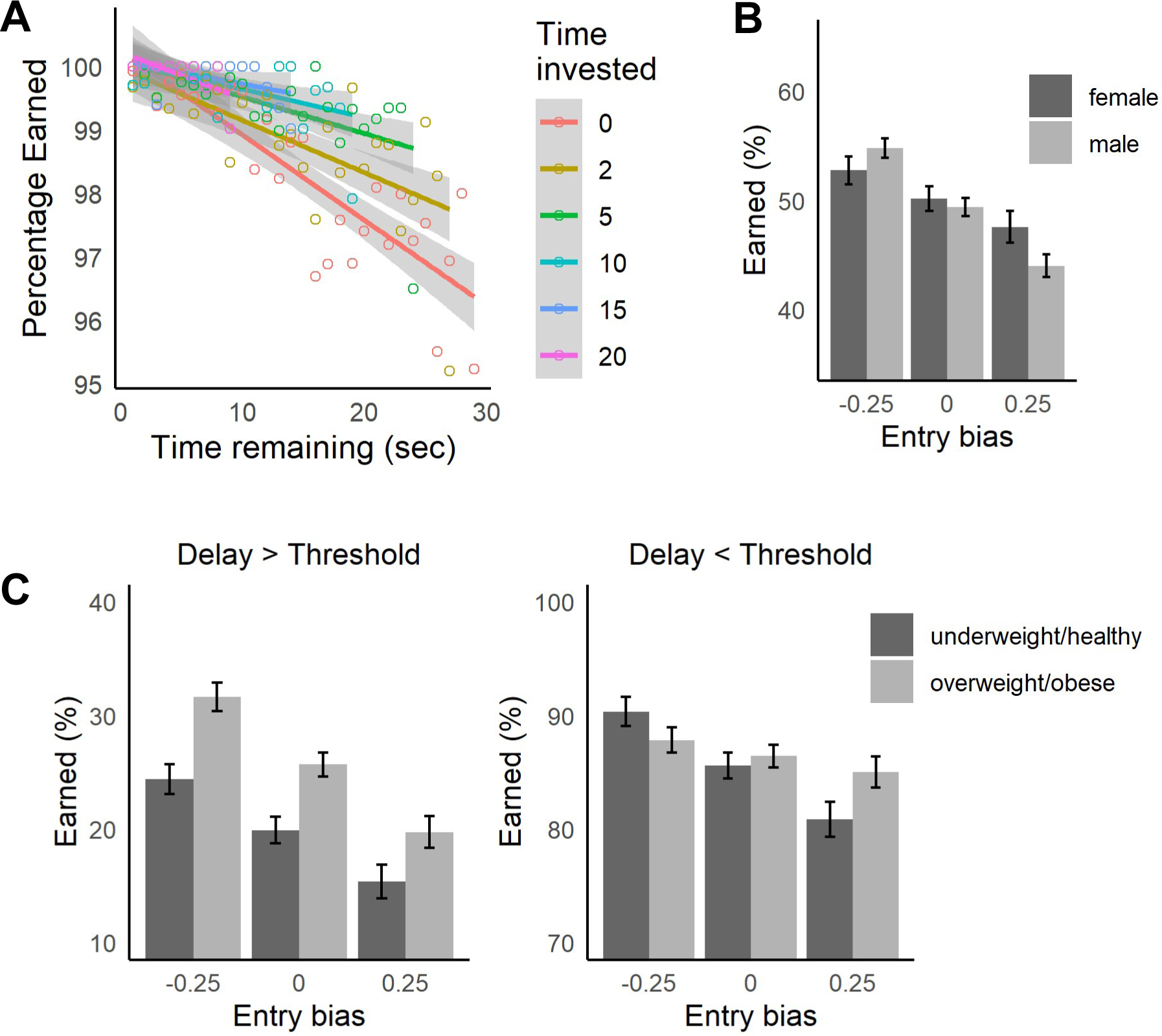
Replication of sunk-cost results from the Restaurant Row and Web-Surf. A: Quit rates (leaving before the delay was completed) increased as a function of delay ( decreased for longer delays (black points and grey area in each plot). The probability of staying for the remainder of the delay increased as a function of amount of time invested (investments of 2-20 seconds shown). Shaded areas indicate the 95% confidence intervals. B: Percentage of rewards earned by gender, estimated for entries to the center (0), left (−0.25) and right (+0.25) of center. Participants were more likely to accept rewards when entering to the left of center, and more likely to skip them when entering to the right of center, with males showing stronger sensitivity to this bias. C: Relationship to BMI. Overweight/obese individuals (BMI ≥ 25) were more likely to accept low-value offers (above threshold) and less sensitive to entry bias for high-value offers (below threshold) compared to underweight/healthy individuals (BMI < 25).

The low quit rates observed across the four versions of these tasks may indicate that by the point at which participants reached the platform and initiated the delay, they had already made a substantial investment, and that their behavior was already influenced by sunk-costs (involved in making a choice). In that case, a better measure of sensitivity to sunk-costs might be observed earlier in the process of deciding to accept or reject an offer. On the Movie and Candy Row, another sunk-cost (besides the amount of time invested in waiting during the delay) can be investigated at the point at which participants receive an offer after entering the offer zone. As participants approached an offer zone, no constraints were placed on the position of the participants within the width of the hallway leading into the offer zone. On many occasions, participants entered with a bias to the left (towards the waiting zone) or right (towards the exit) of the center of the hallway (for example, see Figure 6A where the participant shows a bias towards the waiting zone). Based on the average movement speed, approximately 1.25 seconds were required to cross the full width of the hallway. As the median decision latency (calculated within, then across participants) to make a choice when participants entered near the center of the hallway was 2.7 seconds, a substantial bias in the location where participants entered the offer zone (towards the left or right of center) substantially impacted the amount of time required to make a choice. The location that a participant enters the offer zone can thus be considered a sunk-cost in that participants made their entry before knowing what the quality of an offer they would receive, and the cost of their subsequent decision was strongly impacted by their entry bias.

**Figure 6.**
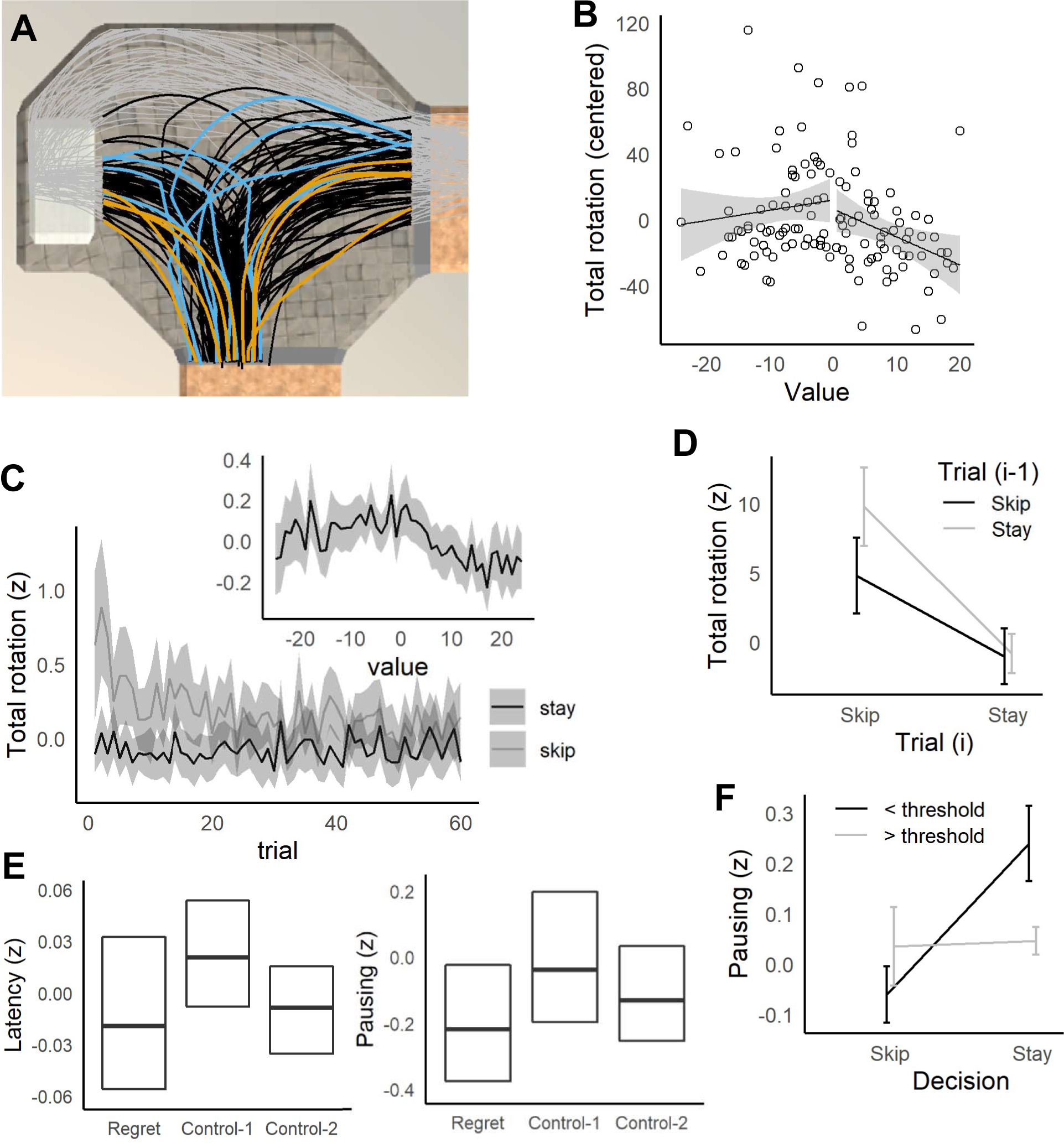
Offer zone behavior. A: Example of position data from one participant (MRM336), rotated to overlay a single offer zone. All position data are shown as grey points, black lines indicate paths taken up to the point where the participant left the offer zone. Randomly selected trials when the participant’s choice time was slow (blue) versus fast (orange) show that the participant tended to task longer paths and make corrections on slow trials. B: Relative to offer value (threshold – offer), total angular rotation was elevated for difficult offers (value = 0) for MRM336. C: Across participants, total rotation was elevated for difficult decisions (value = 0). Total rotation decreased across trials for skip decisions, but not stay decisions. E: Participants made faster decisions on regret trials, and were less likely to pause, compared to control-1 trials. F: Participants spent more time paused when making decisions that were not consistent with their delay thresholds (skipping offers that were < threshold, and accepting offers > threshold).

To assess the impact of entry bias on stay/skip decisions, a linear mixed-effects model was fit with Choice (1= stay, 0 = skip) as the dependent variable, value (coded dichotomously as 0 = < threshold, 1 = > threshold) and entry bias (−50% to +50% of the hallway width), gender, and the interactions of these factors as fixed-effect independent variables and participant as a random effect (*Choice ∼ value type + entry bias + gender + value type:entry bias + value type:gender + entry bias:gender + value type:entry bias:gender + (1|participant)*). As expected, participants were more likely to accept high value offers (delay < threshold) compared to low value offers (delay > threshold, see Supplementary Table 3, p < 0.001, p-adj = 0.002). The probability of accepting an offer also declined as the entry bias increased, and participants entered the offer zone to their right (p < 0.001, p-adj = 0.002). There was also a significant *entry bias:gender* interaction, where males showed a stronger sensitivity to entry bias overall (Figure 5B). This gender difference may also represent a confound with age, as when the data used for the model was restricted to participants 40 years old or younger, the interaction was weakened, though the uncorrected p-value was still significant (β = 0.092, 95% CI = 0.012, 0.17, p = 0.034, p-adj = 0.068). Across gender, when comparing estimates of the percentage of rewards earned based on this model (using percentage of rewards earned as the dependent) for an entry bias near 0 (entering at the center of the hallway) to when participants were biased towards the left by 25% (halfway between the center of the path and the waiting zone), the percentage of poor offers accepted increased 5.5%. And, when participants were biased towards the right by 25% (towards the exit), the percentage of good offers skipped was 3.7% lower. These data support the interpretation that entry bias represents a sunk-cost, and that participants were more likely to reject good offers and accept bad offers if they had invested effort in moving towards the disadvantageous choice (towards skipping on trials with good offers, and towards staying on trials with bad offers) before the value of the offer was revealed.

The relationship of choices to entry bias was also sensitive to BMI, but not to smoking status in these samples. When BMI group (< or ≥ 25) was used in the model instead of gender, there was a significant *value type:entry bias:bmi group* interaction (Supplementary Table 3, p < 0.001, p-adj = 0.002). Separate models were conducted for low and high value offers, to investigate the 3-way interaction. For low value offers (delay > threshold), overweight/obese individuals were more likely to accept an offer overall (Figure 5C, *bmi group*: β = −0.050, 95% CI = −0.087, −0.015, p = 0.002, p-adj = 0.003), but the interaction of entry bias with BMI group was not significant (β = 0.008, 95% CI = −0.036, 0.059, p = 0.74, p-adj = 0.74). For high value offers (delay < threshold), there was a significant *entry bias:bmi group* interaction (β = −0.054, 95% CI = −0.096, −0.014, p = 0.010, p-adj = 0.013), where overweight/obese individuals were less sensitive to entry bias than underweight/healthy individuals. In a model where both gender and BMI group were included, along with all interactions of the factors, the same basic pattern of results was obtained, however, the 4-way interaction of *value type:entry bias:gender:bmi group* approached significance (β = 0.055, 95% CI = 0.0005, 0.16, p = 0.054, p-adj = 0.12). However, when the sample was restricted to participants who were 40 years old and younger, this interaction was reduced (p = 0.92, p-adj = 0.92), indicating a potential confound with age.

Smoking status was not well related to sensitivity to entry bias. When smoking status (non-smoker, smoker) was used in the model instead of gender, the effect of value type, entry bias and their interaction remained significant (Supplementary Table 3, ps < 0.025, p-adjs < 0.048).

### Overriding preferences

Sweis and colleagues (2018) have reported that time spent in the offer zone and VTE are increased when mice skip offers for their most preferred food, and that increased VTE is associated with improved decision-making (associated with an increased probability of skipping low-value offers, which are above the threshold for the restaurant). To determine if similar patterns were obtained for humans, we examined decision latency and offer zone behaviors when participants made decisions (stay/skip) which were consistent or inconsistent with their preferences.

In contrast to results on the Restaurant Row with mice, humans tested on the Movie/Candy Row tasks did not show increased decision times when skipping offers for their most-preferred rewards (Supplementary Figure 3). In an ANOVA with post-task Ranking (1-4) and Decision (skip/stay) as within-subjects factors, and Gender (female/male) as a between-subjects factor, there was no evidence for a Rank × Decision interaction (F(3, 483) < 1, n.s., η^2^_p_ = 0.003) for decision latencies (z-scored). There was a significant main effect of Rank (F(3, 483) = 2.9, p = 0.034, η^2^_p_ = 0.018), with participants making faster choices in their most-preferred reward location. There was a significant Gender × Decision interaction (F(1, 161) = 9.5, p = 0.003, η^2^_p_ = 0.056), where males had longer decision latencies when skipping offers, and females having longer decision latencies when accepting offers. No other main effects or interactions were significant (p > 0.19).

While we did not find that skip decisions were significantly slower for preferred rewards, decision latencies were elevated on trials in which participants violated their delay thresholds. Decision latencies (z-scored) were higher when accepting low-value offers (above threshold) or rejecting high-value offers (below threshold, see Supplementary Figure 2). Analyzing decision latencies (z-scored) using an ANOVA with Gender as a between-subjects factor, and Decision (skip/stay) and Value (< threshold, > threshold) as within-subjects factor, there was a significant Value × Decision interaction (F(1, 211) = 32.9, p < 0.001, η^2^_p_ = 0.13). As with the analysis for rank, there was a significant Gender × Decision interaction (F(1, 211) = 8.5, p < 0.001, η^2^_p_ = 0.039), and the Gender × Value × Decision interaction approached significance (F(1, 211) = 3.9, p = 0.05, η^2^_p_ = 0.018). Examining females and males with separate ANOVAs, there a significant Value × Decision interaction for both genders (p < 0.03), and the main difference observed was that females did not show increased latencies when accepting low-quality offers. This gender difference may indicate a possible confound with age: more of the younger participants were male undergraduates. Overall, the average difference in decision latencies between choices that were inconsistent versus consistent with the delay threshold was stronger for younger male and female participants (Supplementary Figure 1B), and it is possible that a factor relevant to decision latencies (such as video game experience) was related to age. When the original ANOVA was restricted to participants who were 40 years and younger, the Gender × Value × Decision interaction was no longer significant (p = 0.28), while the significant Value × Decision interaction remained (p < 0.001).

These results contrast with results in mice (Sweis et al., 2018), where choices were fastest when accepting offers in general (both above and below threshold), indicating a possible species difference, similar to that observed for regret. When successfully rejecting offers above threshold, mice were slower to make a decision and engaged in more VTE, suggesting a specific role for deliberation in rejecting low-quality offers for preferred rewards. In contrast, humans tested here in the virtual version of the task made their fastest choices when acting consistently with their preferences (accepting high-value offers, rejecting low-value offers), and were slower when acting inconsistently with these preferences.

Several other behavioral measures were also sensitive to these delay threshold violations, and were elevated when participants made decisions that were not consistent with their delay thresholds. In additional ANOVAs conducted for each behavioral measure, there were significant Rank × Decision interactions for total distance travelled (z-scored, F(1, 211) = 10.3, p = 0.002, η^2^_p_ = 0.0.047), total rotation (mean-centered, F(1, 211) = 10.4, p = 0.002, η^2^_p_ = 0.047), rotation reversals (F(1, 210) = 13.5, p < 0.001, η^2^_p_ = 0.06), pausing (z-scored, F(1, 211) = 19.3, p < 0.001, η^2^_p_ = 0.084), as well as reaction time (F(1,170) = 4.7, p = 0.031, η^2^_p_ = 0.027). The results for total rotation and rotation reversals were qualified by significant Gender × Value × Decision interactions (total rotation: F(1, 211) = 10.5, p = 0.001, η^2^_p_ = 0.047, reversals: F(1, 210) = 12.2, p < 0.001, η^2^_p_ =0.055), and separate ANOVAs by gender indicated that Value × Decision interactions were significant in males (ps < 0.001), but not in females (ps > 0.85), and this pattern of results was not changed by restricting the analysis to participants 40 years and younger.

These results indicate that when participants made decisions inconsistent with their delay thresholds, their latency to make a choice was lengthened, during which they travelled farther and tended to pause for a longer duration. Males were also more likely to change their direction of rotation, and rotate more overall before completing their choice on these trials.

### Sequential choices

Abram and colleagues (2019) also have found that humans tested on the Web-Surf were slowest to make a decision when skipping a trial after accepting the previous offer. Similar results were obtained in the Movie/Candy Row tasks, with decision latencies significantly higher on trials where participants skipped an offer after accepting the previous offer (Figure 4B). In a three-factor ANOVA with decision on the current trial *i* (Decision_(i)_: skip, stay) and decision on previous trial *i-1* (Decision_(i-1)_: skip, stay) as within-subjects factors, and Gender as a between-subjects factor, there a significant main effect of Decision_(i-1)_ (F(1, 242) = 46.1, p < 0.001, η^2^_p_ = 0.024), but not Decision_(i)_ (F(1, 242) = 1.9, p = 0.17, η^2^_p_ = 0.005). Importantly, there was a significant Decision_(i)_ × Decision_(i-1)_ interaction (F(1, 242) = 29.1, p < 0.001, η^2^_p_ = 0.015). Decision latencies were slower when participants skipped an offer after accepting the previous offer (Stay/Skip) compared to Skip/Skip (t(187) = 7.5, p < 0.001), Stay/Stay (t(187) = 4.3, p < 0.001) and Skip/Stay (t(187) = 4.5, p < 0.001) conditions.

However, these results were qualified by a significant Gender × Decision_(i)_ × Decision_(i-1)_ interaction (F(1, 242) = 4.2, p = 0.042, η^2^_p_ = 0.002). Separate ANOVAs by gender indicated that the Decision_(i)_ × Decision_(i-1)_ was stronger in males (F(1, 157) = 45.7, p < 0.001, η^2^_p_ = 0.030) than females (F(1, 85) = 3.5, p = 0.067, η^2^_p_ = 0.007). As with results described above for when participants were overriding their preferences, this apparent gender difference may indicate a confound with age. When analyses were restricted to participants 40 years and younger, the Gender × Decision_(i)_ × Decision_(i-1)_ interaction was not significant (F(1, 199) = 2.3, p = 0.13, η^2^_p_ = 0.001).

On Stay/Skip trials, participants also travelled farther on average, paused longer, and were more likely to reverse their direction of rotation (Supplementary Figure 5) compared to Skip/Skip trials. In separate ANOVAs for other behavioral measures, there were significant Decision_(i)_ × Decision_(i-1)_ interactions for the distance travelled (z-scored, F(1, 242) = 23.3, p < 0.001, η^2^_p_ = 0.014), pausing (F(1, 238) = 8.0, p = 0.005, η^2^_p_ = 0.007), and rotation reversals (F(1, 242) = 44.1, p < 0.001, η^2^_p_ = 0.015). The pattern for total rotation (Figure 6D) was similar to that of distance travelled, but the interaction was not significant (total rotation: F(1, 242) = 2.8, p = 0.39, η^2^_p_ < 0.001). Participants spent more time paused on Stay/Skip trials compared to Skip/Skip trials (t(239) = 5.0, p < 0.001), but overall participants spent more time paused if they accepted the previous offer (main effect of Decision_(i)_: F(1, 238) = 35.6, p < 0.001, η^2^_p_ = 0.054).

Similarly, participants travelled shorter distances (F(1, 239) = 55.4, p < 0.001, η^2^_p_ = 0.11), and rotated less overall F(1, 238) = 12.9, p < 0.001, η^2^_p_ = 0.027) when accepting offers, but these effects were consistent with the overall bias for participants to enter the left-hand side of the offer zone (entry bias: M = −12.6%, SD = 19.3%). No main effects or interactions were significant for reaction time (z-scored, ps > 0.14, η^2^_p_ < 0.0025), except that reaction times were slower if participants had accepted the previous offer (main effect of Decision_(i-1)_: F(1, 238) = 5.7, p = 0.018, η^2^_p_ = 0.009). Except for the initial ANOVA for decision latencies, none of the Gender × Decision_(i)_ × Decision_(i-1)_ interactions were significant (ps > 0.10, η^2^_p_ < 0.0016).

### Deliberation

Rats and mice demonstrate vicarious-trial-and-error (VTE) in the offer zone, when deliberating on which offers to accept (Steiner, Adam P. & Redish, 2014; Sweis et al., 2018; Sweis et al., 2018). Steiner and Redish first demonstrated that VTE (measured as the integrated absolute angular change in the orientation of motion of the head) was sensitive to value (the difference between the delay threshold and the delay offered on a given trial) in rats tested on the Restaurant Row. For difficult offers (value = 0, for delays close to the threshold), rats and mice spend more time in the offer zone and engage in more VTE, consistent with the proposal that VTE is a behavioral correlate of deliberation which is expected to be enhanced for difficult offers. Similarly, humans tested on the Web-Surf take more time to make a stay/skip decision for difficult offers, where the value = 0 (Abram et al., 2019). To examine the relationship between deliberation and behavior in humans further, the relationship between value (delay offered on a trial minus the participant’s delay threshold) and each behavioral measure was examined. We predicted that more difficult offers (value = 0), would require more deliberative processing, and that these behavioral measures (related to the time taken to make a decision, or the movements while in the offer zone) would be sensitive to value, and peak at value = 0.

For decision latency, the pattern expected for deliberation was observed (Figure 4C, inset), with the highest latencies observed for the most difficult offers (peaking as the delay offered approached the participant’s threshold for that reward type). A linear mixed-effects model was fit with decision latency (z-scored) as the dependent variable, absolute value of the offer (|value|: 0 to 25 seconds) as a continuous predictor, and value type (coded dichotomously as 0 = < threshold or 1 = > threshold) and their interactions included as fixed-effect independent variables and participant as a random effect (*z_latency_ ∼ |value| + value type + gender + |value|:value type + |value|:gender + value type:gender + |value|:value type:gender + (1|participant)*). Decision latencies increased significantly for difficult offers, as value approached 0 (Supplementary Table 4, p < 0.001, p-adj = 0.004). No other effects involving |value| or value type were significant (ps > 0.18, p-adj > 0.34).

Similar to the results for decision latencies, separate regressions for other behavioral measures found no significant interactions with gender, except for rotation reversals (*|value|:gender*, and *|value|:value type:gender*, ps < 0.007, p-adjs = 0.008) and reaction time (*|value|:gender*, p < 0.001, p-adj = 0.008). Results are given in Supplementary Table 4 for distance travelled, total rotation, and time spent paused calculated across both genders. Separate regressions were conducted by gender for rotation reversals and reaction time, and results are given in Supplementary Table 5. As shown in Figure 6C and Supplementary Figure 6, when participants were presented with difficult offers (where the delay offered approached the participant’s threshold), participants were not only slower to make decisions but they were also more likely to pause, travel farther, rotate more, and change their direction of rotation (|value|, ps < 0.005, p-adjs < 0.009). For rotation reversals (in males), total rotation, and distance travelled, the sensitivity to |value| differed for low and high value offers (|value|:value type, ps < 0.027, p-adjs < 0.027), with steeper slopes for high value offers for total rotation and distance travelled.

Also, females (but not males) showed some evidence for deliberation before starting their path through the offer zone, with a significant effect of |value| for reaction time (p < 0.001, p-adj = 0.002). This gender difference and the significant effects of gender for rotation reversals may be explained by differences in age (which may be confounded with video game experience) between the males and females in the sample (as was explored above in analyses of sequential choices and when participants were overriding their preferences). When the regressions were restricted to participants who were 40 years old or younger, the *|value|:gender* interaction term for reaction time was no longer significant (p = 0.074, p-adj = 0.28), nor were any interactions with gender for rotation reversals (ps > 0.32, p-adjs > 0.42).

The tendency for decision latencies to be elevated for difficult offers was also related to BMI, but not to smoking status (Supplementary Table 6). Adding smoking status to the regressions for decision latency revealed no significant effects or interaction terms with smoking status (ps > 0.24, p-adjs > 0.47). Adding BMI group (< or ≥ 25) to the decision latency regression model did reveal a significant *|value|:value type:BMI group* interaction term (p = 0.008, p-adj = 0.016). Separate regressions by BMI group found that both groups demonstrated a significant relationship to |value| (ps < 0.001, p-adjs < 0.005). As shown in Figure 4D, for underweight/healthy individuals, the slope for |value| was similar for offers below and above threshold, with a weak trend towards stronger sensitivity for bad offers (*|value|:value type* interaction: β = 0.006, 95% CI = −0.00001, 0.013, p = 0.07, p-adj = 0.07). In contrast, overweight/obese individuals were more sensitive to |value| for good offers compared to bad offers *(|value|:value type* interaction: β = −0.007, 95% CI = −0.014, −0.0008, p = 0.034, p-adj = 0.045).

## Discussion

Behavior on the Movie Row and Candy Row tasks largely replicated published behavioral results from rodents (in the Restaurant Row navigation task) and humans (in the Web-Surf experiential foraging task), demonstrating the utility of this set of tasks to capture information about multiple dimensions of decision-making, including deliberation, regret and sunk-costs. Our results with the Movie Row and Candy Row extend the comparison of human and rodent decision-making, by demonstrating that when faced with difficult offers (near threshold) and when making acting against one’s preferences (skipping high-value offers and accepting low-value offers), humans not only take longer to make a decision, but tend to pause longer, rotate farther, change rotation direction, and travel further. These results demonstrate the existence of vicarious trial-and-error (VTE) in humans which share many similarities to rodent behavior on the Restaurant Row, and suggest that VTE in navigation may be a behavioral correlate of deliberation shared across humans and rodents.

Rats and mice on the Restaurant Row (Steiner, Adam P. & Redish, 2014; Sweis et al., 2018) take more time to decide to accept or skip difficult offers (close to threshold). These observations are consistent with the proposal that deliberative processing is computationally slow (time-intensive), involving a search through potential future states to find the best decisions or actions to take (Redish, 2013). In rodents, difficult decisions on the Restaurant Row task are also associated with an increase in VTE, which has been proposed to be a behavioral correlate of deliberation in rodents (Steiner, Adam P. & Redish, 2014). During VTE in rats, neural activity in the hippocampus represents “sweeps” through potential future trajectories (Johnson & Redish, 2007), while activity in the orbitofrontal cortex and ventral striatum represent potential goal locations (Stott & Redish, 2014).

While decision times in humans are also slower when faced with difficult decisions on the Web-Surf task (Abram et al., 2019), behavioral correlates of deliberation in humans (analogous to VTE) have not been well described. In visuospatial tasks, primates (human and nonhuman) show evidence for a visual search pattern (‘saccade-fixate-saccade’) that is similar to VTE, in which subject’ fixations alternate between targets during difficult decisions (reviewed in Redish, 2016). Revisitation, a return of fixation to the previous stimulus, during study is associated with better subsequent memory for items, and is reduced in amnesiacs with hippocampal damage, similar to results in rodent VTE (Voss et al., 2011). During difficult perceptual discriminations, revisitation is also associated with improved performance and increased hippocampal activity (Voss & Cohen, 2017).

While patterns of eye movements, such as the saccade-fixate-saccade pattern, share a number of properties with rodent VTE in rats, it is unclear if humans and other primates demonstrate similar VTE behaviors during navigation that are directly comparable. In rats, VTE was initially characterized in tasks where animals were presented with discrete choices (such as two alleys in a maze, and trained to make difficult discriminations, Muenzinger & Gentry, 1931). At these choices, VTE was characterized generally as a “hesitating, looking-back-and-forth, sort of behavior which rats can often be observed to indulge in at a choice-point before actually going one way or the other” (Tolman, 1948, p. 196-7). Muenzinger (1938) identified two primary patterns behavior that characterized VTE in rats trained in discrimination tasks on a T-maze: “Our criterion for recording *VTE* behavior in any one trial was a facing into one alley before the other one, whether right or wrong, was entered. This alternation in facing the two choice alleys is accomplished in various ways by white rats. The most common way is for the rat to stop at a mid-point between the alleys and turn his head first towards one and then towards the other alley. But he may also approach the entrance to the alley and orient his whole body towards it and then turn and approach the other alley in a similar way” (p. 77). These two patterns of behavior are captured in studies by Redish and colleagues by the metric IdPhi (the total angular rotation as animals pass through an offer zone, or in a fixed window of time after entering the offer zone, Papale et al., 2012; Steiner, A. P. & Redish, 2012), though when body position coordinates are used rather than head position coordinates, the IdPhi is likely less sensitive to the first type of VTE behavior described by Muenzinger (when animals look back and forth while paused).

In virtual navigation research with humans, one recent study by Santos-Pata and Verschure (2018) found elevated VTE in a virtual navigation task (applying the IdPhi measure to the rotation of the head of the participant’s avatar, and quantifying oscillations in head orientation) early in training and at early, high-cost choice points in a multiple T maze task. In that task, rotation was controlled using a mouse, while movements were controlled using a keyboard (a common key binding for first-person perspective games presented on laptops or desktop computers). The patterns described by Santos-Pata and Verschure are consistent with the proposal that these behaviors represent VTE during the use of hippocampally-dependent place strategies in navigation, but the data do not specifically address if VTE is enhanced in individuals using place-strategies, versus striatally-dependent response-strategies. Our results extend this work by examining a range of VTE behaviors in situations which promote deliberation, and demonstrating that behaviors which are comparable to rodent VTE are enhanced when humans appear to be deliberating between options or when acting against their preferences. Our results seem to be most consistent with the second VTE pattern described by Muenzinger (1938), in that participants who took longer to make a choice were likely to make an initial commitment to one option (to stay or skip), then changing direction one or more times before making a final choice (see examples in Figure 6A). In our study, movement was also controlled simply using keyboard keys to rotate, which may impact the type of VTE that is observed in our task, compared to that of Santos-Pata and Verschure (2018). However, using keyboard keys, rather than using a mouse to control orientation, may be preferable for testing participants with a wide range of computer experience, and our results suggest that even with this simpler set of movement controls, we see robust evidence for VTE in the Movie and Candy Row tasks.

Beyond deliberation, the results from the Movie and Candy Row tasks also replicate and extend previous findings from the Restaurant Row and Web-Surf. On the Movie and Candy Row, participants demonstrated consistent preferences for video types, which agreed well with their stated preferences. In participants tested on both tasks (one using videos as rewards, and the other using actual food rewards), performance was correlated, where individuals who were more willing to wait for video rewards tended to also be willing to wait for palatable foods (candy and snacks). While there may be situations in which food is a preferred reward, these results support the generalizability of research using videos as rewards (in the Web-Surf and Movie Row), an approach that is more suitable for online data collection and the testing of larger samples.

One species difference noted here was during a regret-inducing situation (receiving a low-value offer after skipping a high-value offer), humans were less likely to accept the low-value offer, and appeared less likely to deliberate (spent less time in the offer zone, rotated less, were less likely to pause and tended to travel a shorter distance, compared to control trials). Compared to control conditions, rats in regret-inducing situations are more, rather than less, likely to accept low-value offers (Steiner, Adam P. & Redish, 2014), and are more likely to look back at the previous reward location and consume the rewards earned more quickly. During regret-inducing trials, activity of neural ensembles in the orbitofrontal cortex and ventral striatum also tended to represent the previous location (where rats had skipped a high-value offer), activity which may be important in reevaluating past decisions to guide future behavior. Interesting, neural representations in humans (assessed by fMRI) are enhanced for the current location (rather than the previous one) on regret-inducing trials (Abram et al., 2019), the results presented here further support species differences in behavioral and neural processes associated with regret.

Replicating previous research, participants also demonstrated a sunk-cost bias, with participants’ likelihood of completing a delay increasing as a function of the amount of time already invested in waiting. A separate measure of sunk-costs was also identified, based on the initial position at which participants entered the offer zone (towards the reward waiting zone, or towards the offer zone exit). While participants on the Movie Row and Candy Row tasks very rarely quit after initially accepting an offer (limiting the variability across participants or task experience), the participants’ entry bias represented an investment with a significant impact on the cost for accepting or rejecting an offer, which had a substantial impact on the willingness of a participant to accept an offer.

The Restaurant Row and Web-Surf have also shown promise as tools to understand how decision-making is related to vulnerability to addiction and the impact of drug exposure. Results with cocaine and morphine abstinent mice have shown that these tasks have promise in the study of how drug use and cessation specifically impact decision-making (Sweis et al., 2018). Similarly, in a version of the Web-Surf which incorporates risky offers (Abram et al., 2019), individuals with high trait externalizing (who thus may be at risk for addiction) were less likely than those with low trait externalizing to avoid risky offers after a loss, potentially signaling an impairment in learning from risky losses. These studies indicate the potential for experiential foraging tasks to capture decision-making processes that are relevant to human disorders, such as addiction. To further explore the potential of these tasks, we also explored the relationship of delay-discounting measures and foraging behavior to smoking status and BMI.

Using two measures of delay-discounting (the Monetary and Food Choice Questionnaires, MCQ and FCQ), we found that participants who were less willing to wait for hypothetical monetary rewards (MCQ) were also more less to wait for hypothetical food rewards (FCQ), indicating individual differences in delay-discounting across two types of reinforcer (a primary reinforcer, food, and a secondary reinforcer, money). Similarly, we found that delay thresholds and the proportion of rewards earned on the Movie Row and Candy Row tasks were correlated, indicating an individual difference in willingness to wait for rewards across two reinforcer types (movies and food, both of which are primary reinforcers). However, willingness to wait for hypothetical rewards (in the MCQ and FCQ) were not related to willingness to wait for actual rewards (in the Movie Row and Candy Row), indicating that these two measures assess different dimensions of decision-making. These findings are consistent with results obtained using a version of the Web-Surf that incorporated risk (Abram et al., 2019), where measures of discounting rates (based on the delay until or probability of receiving a large reward) did not relate to trait externalizing, and did not account for the relationship between externalizing and Web-Surf task performance. Our results are also consistent with the findings of studies which compare foraging to forced-choice tasks which provide equivalent rewards. In these cases, behavior observed when animals are given a forced-choice between an immediate small reward and a delayed large reward can deviate strongly from behavior in the same animals when the options are presented as a stay/leave foraging decision (Carter & Redish, 2016; Stephens, 2008).

Consistent with published reports, smokers more steeply discounted hypothetical future monetary rewards compared to non-smokers. And, individuals reporting higher BMI (≥ 25) did have somewhat higher discounting rates (k) for monetary rewards, consistent with a previous report (Jarmolowicz et al., 2014) which found significantly higher discounting (on the MCQ) for overweight/obese compared to underweight/healthy participants. The weaker relationship observed between BMI and discounting that we observed does fit within the larger literature on delay-discounting, with some meta-analyses supporting for a relationship (Amlung, Petker, Jackson, Balodis, & MacKillop, 2016), and others finding mixed evidence (Tang Jianjun, Chrzanowski-Smith, Hutchinson, Kee, & Hunter, 2019), and more so when hypothetical rewards are used (such as in the MCQ and FCQ).

In contrast, on the foraging tasks neither smoking status nor BMI group were strongly related to delay thresholds nor to the proportion of rewards earned. And, the strongest trend observed was that smokers were, if anything, more willing to wait for rewards (with a trend towards longer delay thresholds and earning more rewards). The trend for smokers may have been driven by a tendency for smokers to report higher subjective enjoyment in the Movie Row task (in the post-video star ratings), and thus may have been more willing to wait for videos because they found them more rewarding. Such a result seems consistent with other work demonstrating that, at least for acute administration, nicotine can enhance the rewarding effects of some non-drug rewards (music and video stimuli, but not money, Perkins, Karelitz, & Boldry, 2017).

BMI was related to decision-making, as overweight/obese individuals were more likely to accept poor offers (higher than the participant’s threshold) overall, and also showed reduced sunk-cost bias for good offers (below threshold), as assessed by the relationship between entry bias and stay/skip decisions. Smoking status was not significantly related to sunk-cost sensitivity in this sample, though smokers did show a nonsignificant trend in a similar direction (towards reduced sensitivity to the cost represented by entry bias). While it is unknown if obesity and nicotine use are related to sunk-cost sensitivity using more traditional measures, a study by Fujino and colleagues (2018) found no differences in sunk-cost sensitivity in males with gambling disorders compared to male healthy controls on a task based on a scenario used in Arkes and Blumer’s (1985) study of the sunk cost effect. However, sunk-cost sensitivity in the gambling disorder sample was negatively correlated with duration of abstinence from gambling, suggesting that hypothetical decisions involving sunk-costs may be disrupted during addiction. The relationship between sensitivity to sunk-costs as assessed in foraging tasks (the Web-Surf and Movie/Candy Row) and more standard sunk-cost tasks remains to be determined, and could provide further insight into the ways in which individuals differ in their vulnerability to addictions.

While overweight/obese individuals showed less sensitivity to a measure of sunk-costs for high value offers, their decision latencies also showed stronger sensitivity to value for high value offers, compared to low value offers. Overall, these results suggest that for attractive offers, decisions by high BMI individuals were driven by the value of the offer (and were less impacted by the cost of obtaining the reward).

Research using experiential foraging tasks such as the Restaurant Row and Web-Surf have provided new and important insights into decision-making. Our results with the Movie and Candy Row extend this work: the many cross-species behavioral similarities support the utility of these tasks to not only characterize decision-making systems, but to understand how these systems may contribute to, or be impacted by, a range of behaviors which have profound implications for public health.

## Notes

S.V. Abram is supported by the Department of Veteran Affairs Sierra Pacific Mental Illness Research, Education, and Clinical Center (MIRECC).

## References

1. Abram, S. V., Breton, Y., Schmidt, B., Redish, A. D., & MacDonald III, A. W. (2016). The web-surf task: A translational model of human decision-making. Cognitive, Affective & Behavioral Neuroscience, 16(1), 37–50. doi:10.3758/s13415-015-0379-y

2. Abram, S. V., Hanke, M., Redish, A. D., & MacDonald III, A. W. (2019). Neural signatures underlying deliberation in human foraging decisions. Cognitive, Affective, and Behaviorial Neuroscience

3. Abram, S. V., Redish, A. D., & MacDonald III, A. W. (2019). Learning from loss after risk: Dissociating reward pursuit and reward valuation in a naturalistic foraging task. Frontiers in Psychiatry, 10, 359. doi:10.3389/fpsyt.2019.00359

4. Addicott, M. A., Pearson, J. M., Kaiser, N., Platt, M. L., & McClernon, F. J. (2015). Suboptimal foraging behavior: A new perspective on gambling. Behavioral Neuroscience, 129(5), 656–665. doi:10.1037/bne0000082; 10.1037/bne0000082.supp (Supplemental)

5. Addicott, M. A., Pearson, J. M., Wilson, J., Platt, M. L., & McClernon, F. J. (2013). Smoking and the bandit: A preliminary study of smoker and nonsmoker differences in exploratory behavior measured with a multiarmed bandit task. Experimental and Clinical Psychopharmacology, 21(1), 66–73. doi:10.1037/a0030843; 10.1037/a0030843.supp (Supplemental)

6. Amlung, M., Petker, T., Jackson, J., Balodis, I., & MacKillop, J. (2016). Steep discounting of delayed monetary and food rewards in obesity: A meta-analysis. Psychological Medicine, 46(11), 2423–2434. doi:10.1017/S0033291716000866

7. Arkes, H. R., & Blumer, C. (1985). The psychology of sunk cost. Organizational Behavior and Human Decision Processes, 35(1), 124–140. doi:10.1016/0749-5978(85)90049-4

8. Carter, E. C., & Redish, A. D. (2016). Rats value time differently on equivalent foraging and delay-discounting tasks. Journal of Experimental Psychology: General, 145(9), 1093–1101. doi:10.1037/xge0000196; 10.1037/xge0000196.supp (Supplemental)

9. Emery, R. L., & Levine, M. D. (2017). Questionnaire and behavioral task measures of impulsivity are differentially associated with body mass index: A comprehensive meta-analysis. Psychological Bulletin, 143(8), 868–902. doi:10.1037/bul0000105; 10.1037/bul0000105.supp (Supplemental)

10. Fujino, J., Kawada, R., Tsurumi, K., Takeuchi, H., Murao, T., Takemura, A.,... Takahashi, H. (2018). An fMRI study of decision-making under sunk costs in gambling disorder doi:https://doi.org/10.1016/j.euroneuro.2018.09.006

11. Hendrickson, K. L., Rasmussen, E. B., & Lawyer, S. R. (2015). Measurement and validation of measures for impulsive food choice across obese and healthy-weight individuals. Appetite, 90, 254–263. doi:10.1016/j.appet.2015.03.015 [doi]

12. Jarmolowicz, D. P., Cherry, J. B., Reed, D. D., Bruce, J. M., Crespi, J. M., Lusk, J. L., & Bruce, A. S. (2014). Robust relation between temporal discounting rates and body mass. Appetite, 78, 63–67. doi:10.1016/j.appet.2014.02.013

13. Johnson, A., & Redish, A. D. (2007). Neural ensembles in CA3 transiently encode paths forward of the animal at a decision point. Journal of Neuroscience, 27(45), 12176–12189. doi:10.1523/JNEUROSCI.3761-07.2007

14. Kirby, K. N., Petry, N. M., & Bickel, W. K. (1999). Heroin addicts have higher discount rates for delayed rewards than non-drug-using controls. Journal of Experimental Psychology: General, 128(1), 78–87. doi:10.1037/0096-3445.128.1.78

15. MacKillop, J., Amlung, M. T., Few, L. R., Ray, L. A., Sweet, L. H., & Munafò, M. R., (2011). Delayed reward discounting and addictive behavior: A meta-analysis. Psychopharmacology, 216(3), 305–321. doi:10.1007/s00213-011-2229-0

16. Muenzinger, K. F. (1938). Vicarious trial and error at a point of choice: I A general survey of its relation to learning efficiency Clark University. doi:10.1080/08856559.1938.10533799

17. Muenzinger, K. F., & Gentry, E. (1931). Tone discrimination in white rats Williams & Wilkins Company. doi:10.1037/h0072238

18. Naude, J., Dongelmans, M., & Faure, P. (2015). Nicotinic alteration of decision-making. Neuropharmacology, 96(Pt B), 244–254. doi:10.1016/j.neuropharm.2014.11.021<otherinfo> [doi]</otherinfo>

19. Papale, A. E., Stott, J. J., Powell, N. J., Regier, P. S., & Redish, A. D. (2012). Interactions between deliberation and delay-discounting in rats. Cognitive, Affective & Behavioral Neuroscience, 12(3), 513–526. doi:10.3758/s13415-012-0097-7

20. Perkins, K. A., Karelitz, J. L., & Boldry, M. C. (2017). Nicotine acutely enhances reinforcement from non-drug rewards in humans. Frontiers in Psychiatry, 8, 65. doi:10.3389/fpsyt.2017.00065

21. Redish, A. D. (2016). Vicarious trial and error. Nature Reviews.Neuroscience, 17(3), 147–59. doi:10.1038/nrn.2015.30

22. Redish, A. D. (2013). The mind within the brain: How we make decisions and how those decisions go wrong. New York, NY: Oxford University Press. Retrieved from https://login.wcproxy.palni.edu/login?urlhttp://search.ebscohost.com/login.aspx?direct=true&db=psyh&AN=2013-24085-000&site=ehost-live&scope=sitehttps://login.wcproxy.palni.edu/login?urlhttp://search.ebscohost.com/login.aspx?direct=true&db=psyh&AN=2013-24085-000&site=ehost-live&scope=site

23. Santos-Pata, D., & Verschure, Paul F. M. J. (2018). Human vicarious trial and error is predictive of spatial navigation performance. Frontiers in Behavioral Neuroscience, *12*

24. Steiner, A. P., & Redish, A. D. (2012). The road not taken: Neural correlates of decision making in orbitofrontal cortex. Frontiers in Neuroscience, 6, 131. doi:10.3389/fnins.2012.00131 [doi]

25. Steiner, A. P., & Redish, A. D. (2014). Behavioral and neurophysiological correlates of regret in rat decision-making on a neuroeconomic task. Nature Neuroscience, 17(7), 995–1002. doi:10.1038/nn.3740

26. Stephens, D. W. (2008). Decision ecology: Foraging and the ecology of animal decision making. Cognitive, Affective & Behavioral Neuroscience, 8(4), 475–484. doi:10.3758/CABN.8.4.475

27. Stott, J. J., & Redish, A. D. (2014). A functional difference in information processing between orbitofrontal cortex and ventral striatum during decision-making behaviour. *Philosophical Transactions of the Royal Society of London.Series B*, Biological Sciences, 369(1655) doi:10.1098/rstb.2013.0472

28. Sweis, B. M., Abram, S. V., Schmidt, B. J., Seeland, K. D., MacDonald III, A. W., Thomas, M. J., & Redish, A. D. (2018). Sensitivity to “sunk costs” in mice, rats, and humans. *Science (New York*, N.Y.), 361(6398), 178–181. doi:10.1126/science.aar8644

29. Sweis, B. M., Larson, E. B., Redish, A. D., & Thomas, M. J. (2018). Altering gain of the infralimbic-to-accumbens shell circuit alters economically dissociable decision-making algorithms. PNAS Proceedings of the National Academy of Sciences of the United States of America, 115(27), E6347–E6355. doi:10.1073/pnas.1803084115

30. Sweis, B. M., Redish, A. D., & Thomas, M. J. (2018). Prolonged abstinence from cocaine or morphine disrupts separable valuations during decision conflict. Nature Communications, 9(1), 2521. doi:10.1038/s41467-018-04967-2

31. Sweis, B. M., Thomas, M. J., & Redish, A. D. (2018). Mice learn to avoid regret. Plos Biology, 16(6), e2005853. doi:10.1371/journal.pbio.2005853

32. Tang Jianjun, J., Chrzanowski-Smith, O. J., Hutchinson, G., Kee, F., & Hunter, R. F. (2019). Relationship between monetary delay discounting and obesity: A systematic review and meta-regression. International Journal of Obesity, 43(6), 1135–1146.

33. Tolman, E. C. (1948). Cognitive maps in rats and men American Psychological Association. doi:10.1037/h0061626

34. van der Meer, M. A., Johnson, A., Schmitzer-Torbert, N. C., & Redish, A. D. (2010). Triple dissociation of information processing in dorsal striatum, ventral striatum, and hippocampus on a learned spatial decision task. Neuron-Cambridge Ma*-*, 67(1), 25–32. Retrieved from https://wabash.on.worldcat.org/oclc/649786044

35. Voss, J. L., & Cohen, N. J. (2017). Hippocampal-cortical contributions to strategic exploration during perceptual discrimination. Hippocampus, 27(6), 642–652.

36. Voss, J. L., Warren, D. E., Gonsalves, B. D., Federmeier, K. D., Tranel, D., & Cohen, N. J. (2011). Spontaneous revisitation during visual exploration as a link among strategic behavior, learning, and the hippocampus. Proceedings of the National Academy of Sciences of the United States of America, 108(31), 12581–12582. Retrieved from https://wabash.on.worldcat.org/oclc/5553699464

37. Wang, X. T., Reed, R. N., Baugh, L. A., & Fercho, K. A. (2018). Resource forecasting: Differential effects of glucose taste and ingestion on delay discounting and self-control. Appetite, 121, 101–110. doi:10.1016/j.appet.2017.11.083

