## Supplementary Online Material for "Vicarious trial-and-error is enhanced during deliberation in human virtual navigation in a translational neuroeconomic task"

*Movie Row*

Demonstration versions of the current version of the Movie Row are available at http://virtualnavigationtools.com/WebGL/vns?mrdemowebsf (using the original Web-Surf movies: kittens, bike accidents, dancing, and landscapes) and http://virtualnavigationtools.com/WebGL/vns?mrdemonew (using the second movie set of high-resolution movies: puppies, social interactions, food, and landscapes). This version is currently equivalent to version 4 described in these results, but lacking the fixation task.

*Participants by version*

*Version 1*. Male undergraduates (n = 30), were recruited for a study of the effect of glucose ingestion on decision-making. Participants were asked to fast for at least four hours before the study, and were tested at 8am or 4:30pm each day. Before completing the task, participants provided a blood glucose measurement, completed a version of the Monetary Choice Questionnaire (Using the values and delays, and a related version, Wang, Reed, Baugh, & Fercho, 2018), consumed a drink (water, glucose [26.2g in 12 oz], water after rinsing with 6 oz of the glucose solution), waiting a total of 13 minutes for absorption, and then completing a second blood glucose measurement and an alternate version of the MCQ. Participants then completed the Movie Row task, tested on PCs divided by partitions. Headphones were provided, so that participants could hear the audio presented with each video.

*Version 2*. Male undergraduates (n = 10) and workers from Amazon's Mechanical Turk (mTurk) service (n = 50, 29 females) were recruited to complete the Movie Row task administered online. Participants recruited from mTurk had previously participated in an unrelated virtual navigation study, and completed a screening survey as part of that study.

Participants completed a version of the MCQ as part of the screening questionnaire or immediately before being directed to the Movie Row task.

*Version 3.* Male undergraduates (n = 21) and mTurk workers (n = 97, 51 females, 45 males, 1 non-binary). Undergraduates were tested as a second sample in the glucose study described for version 1 above. Participants from mTurk were recruited for a project examining the relationship of smoking status to delay discounting and performance on the Movie Row task. Participants who reported use of other drugs (besides nicotine or alcohol) in the past 30 days were excluded, as were those who indicated drinking more than 9 times in the previous 30 days, or any alcohol consumption if they also indicated any history of alcohol abuse. From an initial sample of 450 participants who completed the screening questionnaire, 51 smokers and 237 non-smokers were eligible and recruited for the Movie Row task, and a total of 97 participants (20 smokers) completed the Movie Row task.

*Version 4 (Movie Row and Candy Row).* Male undergraduates (n = 83) and members of the local community (n = 34, 19 females, 15 males) completed the Movie Row task. Thirty-one of the male undergraduates completed one session of the Movie Row task online (using the alternate videos, with puppies, social interactions, food, and landscapes). Sixty-nine of the participants also completed the Candy Row task in addition to at least one Movie Row session.

Supplementary Table 1.

*Description of samples tested with each Movie Row version.*

| Version | Location | Source | Videos | n | Age |
|---|---|---|---|---|---|
| 1 | Lab | Undergraduate | original | 30 (0 females) | |
| 2 | Online | Undergraduate | original | 10 (0 females) | |
| | Online | mTurk | original | 50 (29 females) | 43.3 (6.5) |
| 3 | Lab | Undergraduate | original | 21 (0 females) | |
| | Online | mTurk | original | 97 (51 females, 1 non-binary) | 36.4 (10.2) |
| 4 | Lab | Undergraduate | original | 57 (0 females) | 19.1 (1.2) |
| | Online | Undergraduate | new | 30 (0 females) | 18.9 (1.1) |
| | Lab | Community | new | 34 (19 females) | 46.4 (14.1) |

Age and other demographics were not collected for two of the undergraduate samples, but ages were likely to have fallen in the 18-22 age range. Participants were tested with the original Web-Surf videos (kittens, bike accidents, dancing, and landscapes) or a newer, higher-resolution set (puppies, social interactions, food, and landscapes). For version 4, four undergraduate participants completed two versions of the Movie Row task (with different sets of movie clips), with one session completed in the laboratory, and one completed online.

Supplementary Table 2.

*Description of samples by smoking status or BMI group, separated by task and reward type.*

| Task | Videos | Group | n | Age |
|---|---|---|---|---|
| Movie Row | original | Non-smokers | 111 (62 females) | 38.4 (9.9) |
| | | Smokers | 34 (18 females) | 39.9 (9.1) |
| Movie Row | new | Non-smokers | 26 (14 females) | 48.4 (14.7) |
| | | Smokers | 8 (5 females) | 40.0 (9.7) |
| Candy Row | - | Non-smokers | 26 (13 females) | 48.0 (14.2) |
| | | Smokers | 7 (5 females) | 41.4 (9.5) |
| Movie Row | original | Underweight/Healthy | 88 (34 females) | 32.0 (12.3) |
| | | Overweight/Obese | 105 (40 females) | 33.6 (12.0) |
| Movie Row | new | Underweight/Healthy | 17 (9 females) | 37.2 (18.6) |
| | | Overweight/Obese | 27 (10 females) | 42.0 (16.0) |
| Candy Row | - | Underweight/Healthy | 28 (9 females) | 30.2 (17.0) |
| | | Overweight/Obese | 40 (9 females) | 34.2 (16.5) |

Self-reported smoking status and height/weight (used to calculate BMI) were available from a subset of participants tested on the Movie and Candy Row tasks. Smoking status: Non-smokers – reported no current use of nicotine products, Smokers – reported current use of cigarettes or e-cigarettes (vaping). BMI: Healthy/Underweight – BMI < 25.0, Overweight/Obese – BMI $\geq$ 25.0.

Supplementary Table 3.

*Relationship between entry bias and stay/skip decisions*

| | β | 95% CI | p | p-adj |
|---|---|---|---|---|
| Choice (skip/stay) | | | | |
| value type | 0.29 | [0.31, 0.32] | **<0.001** | **0.003** |
| entry bias | -0.22 | [-0.27, -0.18] | **<0.001** | **0.003** |
| gender | -0.016 | [-0.047, 0.016] | 0.35 | 0.35 |
| value type:entry bias | 0.017 | [-0.017, 0.053] | 0.35 | 0.35 |
| value type:gender | -0.015 | [-0.030, -0.002] | **0.036** | 0.057 |
| entry bias:gender | 0.095 | [0.014, 0.17] | **0.016** | **0.032** |
| value type:entry bias:gender | 0.045 | [-0.027, 0.12] | 0.24 | 0.37 |
| *BMI group* | | | | |
| value type | 0.30 | [0.29, 0.31] | **<0.001** | **0.002** |
| entry bias | -0.17 | [-0.21, -0.14] | **<0.001** | **0.002** |
| bmi group | -0.013 | [-0.028, 0.003] | 0.092 | 0.12 |
| value type:entry bias | 0.022 | [-0.011, 0.055] | 0.20 | 0.20 |
| value type:bmi group | 0.011 | [0.005, 0.018] | **<0.001** | **0.002** |
| entry bias:bmi group | -0.030 | [-0.066, 0.008] | 0.12 | 0.13 |
| value type:entry bias:bmi group | -0.064 | [-0.099, -0.032] | **0.002** | **0.003** |
| *Smoking status* | | | | |
| value type | 0.30 | [0.29, 0.032] | **<0.001** | **0.004** |
| entry bias | -0.11 | [-0.18, -0.011] | **0.014** | **0.037** |
| smoker | -0.031 | [-0.071, 0.003] | 0.090 | 0.14 |
| value type:entry bias | 0.084 | [0.015, 0.15] | **0.024** | **0.048** |
| value type:smoker | 0.012 | [-0.005, 0.028] | 0.16 | 0.21 |
| entry bias:smoker | -0.051 | [-0.15, -0.042] | 0.27 | 0.27 |
| value type:entry bias:smoker | -0.051 | [-0.13, 0.026] | 0.21 | 0.24 |

Supplementary Table 3 (continued).

Decisions (0 = skip, 1 = stay) were regressed onto value type of the offer (0 = < threshold, 1 = > threshold), entry bias (-50% to +50% of the hallway width, 0 = hallway center) and the *|value|:value type* interaction (*y ~ value type + entry bias + value type:entry bias*). In separate regressions, smoking status (0 = non-smoker, 1 = smoker) or BMI group (1 = BMI < 25, 2 = BMI ≥ 25) were added to the model along with all interactions between the three terms. Bold indicates $p < 0.05$.

Supplementary Table 4.

*Value models for latency, rotation, distance and pausing*

| | β | 95% CI | p | p-adj |
|---|---|---|---|---|
| Decision latency (z-scored) | | | | |
| \|value\| | -0.012 | [-0.016, -0.008] | **<0.001** | **0.004** |
| value type | -0.037 | [-0.10, 0.022] | 0.22 | 0.34 |
| gender | -0.071 | [-0.14, 0.004] | **0.046** | 0.12 |
| \|value\|:value type | -0.004 | [-0.008, 0.002] | 0.18 | 0.34 |
| \|value\|:gender | 0.002 | [-0.003, 0.008] | 0.47 | 0.47 |
| value type:gender | 0.054 | [-0.062, 0.15] | 0.33 | 0.38 |
| \|value\|:value type:gender | 0.005 | [-0.003, 0.013] | 0.25 | 0.34 |
| Total rotation (z-scored) | | | | |
| \|value\| | -0.011 | [-0.013, -0.009] | **<0.001** | **0.002** |
| value type | 0.015 | [-0.011, 0.038] | 0.33 | 0.25 |
| \|value\|:value type | -0.003 | [-0.005, -0.0007] | **0.006** | **0.008** |
| Distance (z-scored) | | | | |
| \|value\| | -0.008 | [-0.010, -0.006] | **<0.001** | **0.001** |
| value type | -0.047 | [-0.072, -0.023] | **<0.001** | **0.001** |
| \|value\|:value type | -0.002 | [-0.004, -0.0006] | **0.026** | **0.026** |
| Pausing (z-scored) | | | | |
| \|value\| | -0.004 | [-0.006, -0.002] | **<0.001** | **0.004** |
| value type | 0.015 | [-0.009, 0.040] | 0.27 | 0.36 |
| \|value\|:value type | 0.0008 | [-0.001, 0.003] | 0.44 | 0.44 |

Behavioral measures (decision latency, total rotation, distance travelled, and time spent paused) were regressed separately onto absolute value of the offer (|value|), value type (0 = < threshold, 1 = > threshold) and the *|value|:value type* interaction (*y* ~ |*value*| + *value type* + |*value*|*:value type*). Model for decision latencies (described in the main text) includes gender (1 = female, 2 = male) and interactions of |value| and value type with gender (*y* ~ |*value*| + *value type* + *gender* + |*value*|*:value type* + |*value*|*:gender* + *value type:gender* + |*value*|*:value type:gender*). Bold indicates p < 0.05.

Supplementary Table 5.

*Value models by gender for rotation reversals and reaction time*

| | β | 95% CI | p | p-adj |
|---|---|---|---|---|
| Rotation reversals | | | | |
| *Females* | | | | |
| \|value\| | -0.004 | [-0.006, -0.002] | **0.004** | **0.008** |
| value type | 0.010 | [-0.016, 0.039] | 0.48 | 0.62 |
| \|value\|:value type | 0.0006 | [0.002, 0.003] | 0.62 | 0.62 |
| *Males* | | | | |
| \|value\| | -0.010 | [-0.012, -0.008] | **<0.001** | **0.001** |
| value type | 0.017 | [-0.008, 0.038] | 0.14 | 0.14 |
| \|value\|:value type | -0.004 | [-0.006, -0.002] | **<0.001** | **0.001** |
| Reaction time (z-scored) | | | | |
| *Females* | | | | |
| \|value\| | -0.010 | [-0.014, -0.006] | **<0.001** | **0.002** |
| value type | 0.004 | [-0.049, 0.055] | 0.86 | 0.86 |
| \|value\|:value type | 0.002 | [-0.003, 0.006] | 0.41 | 0.55 |
| *Males* | | | | |
| \|value\| | -0.002 | [-0.005, 0.007] | 0.20 | 0.45 |
| value type | -0.010 | [-0.044, 0.026] | 0.58 | 0.58 |
| \|value\|:value type | 0.002 | [-0.001, 0.005] | 0.27 | 0.45 |

Behavioral measures (rotation reversals and reaction times) were regressed separately onto absolute value of the offer (|value|), value type (0 = < threshold, 1 = > threshold) and the *|value|:value type* interaction (*y* ~ *|value|* + *value type* + *|value|:value type*). Separate regressions were performed for females and males. Bold indicates p < 0.05.

Supplementary Table 6.

*Value models by smoking status and BMI group*

| | β | 95% CI | p | p-adj |
|---|---|---|---|---|
| Decision latency (z-scored) | | | | |
| *Smoking status* | | | | |
| \|value\| | -0.013 | [-0.020, -0.006] | **<0.001** | **0.004** |
| value type | -0.086 | [-0.21, 0.05] | 0.20 | 0.47 |
| smoker | -0.046 | [-0.15, 0.061] | 0.38 | 0.61 |
| \|value\|:value type | 0.001 | [-0.010, 0.011] | 0.84 | 0.84 |
| \|value\|:smoker | 0.002 | [-0.006, 0.011] | 0.63 | 0.83 |
| value type:smoker | 0.089 | [-0.050, 0.25] | 0.24 | 0.47 |
| \|value\|:value type:smoker | -0.002 | [-0.013, 0.011] | 0.76 | 0.84 |
| *BMI group* | | | | |
| \|value\| | -0.011 | [-0.014, -0.008] | **<0.001** | **0.004** |
| value type | -0.019 | [-0.078, 0.040] | 0.58 | 0.66 |
| bmi group | 0.033 | [-0.009, 0.073] | 0.11 | 0.14 |
| \|value\|:value type | -0.0003 | [-0.005, 0.004] | 0.91 | 0.91 |
| \|value\|:bmi group | -0.003 | [-0.006, 0.0002] | 0.07 | 0.11 |
| value type:bmi group | -0.080 | [-0.14, -0.019] | **0.008** | **0.016** |
| \|value\|:value type:bmi group | 0.007 | [0.002, 0.011] | **0.006** | **0.016** |

Decision latency ($log_{10}$ transformed and z-scored) was regressed separately onto absolute value of the offer (|value|), value type (0 = < threshold, 1 = > threshold), and smoking status (0 = non-smoker, 1 = smoker) or BMI group (1 = BMI < 25, 2 = BMI $\geq$ 25). All interactions between the three terms were included in the model. Bold indicates p < 0.05.

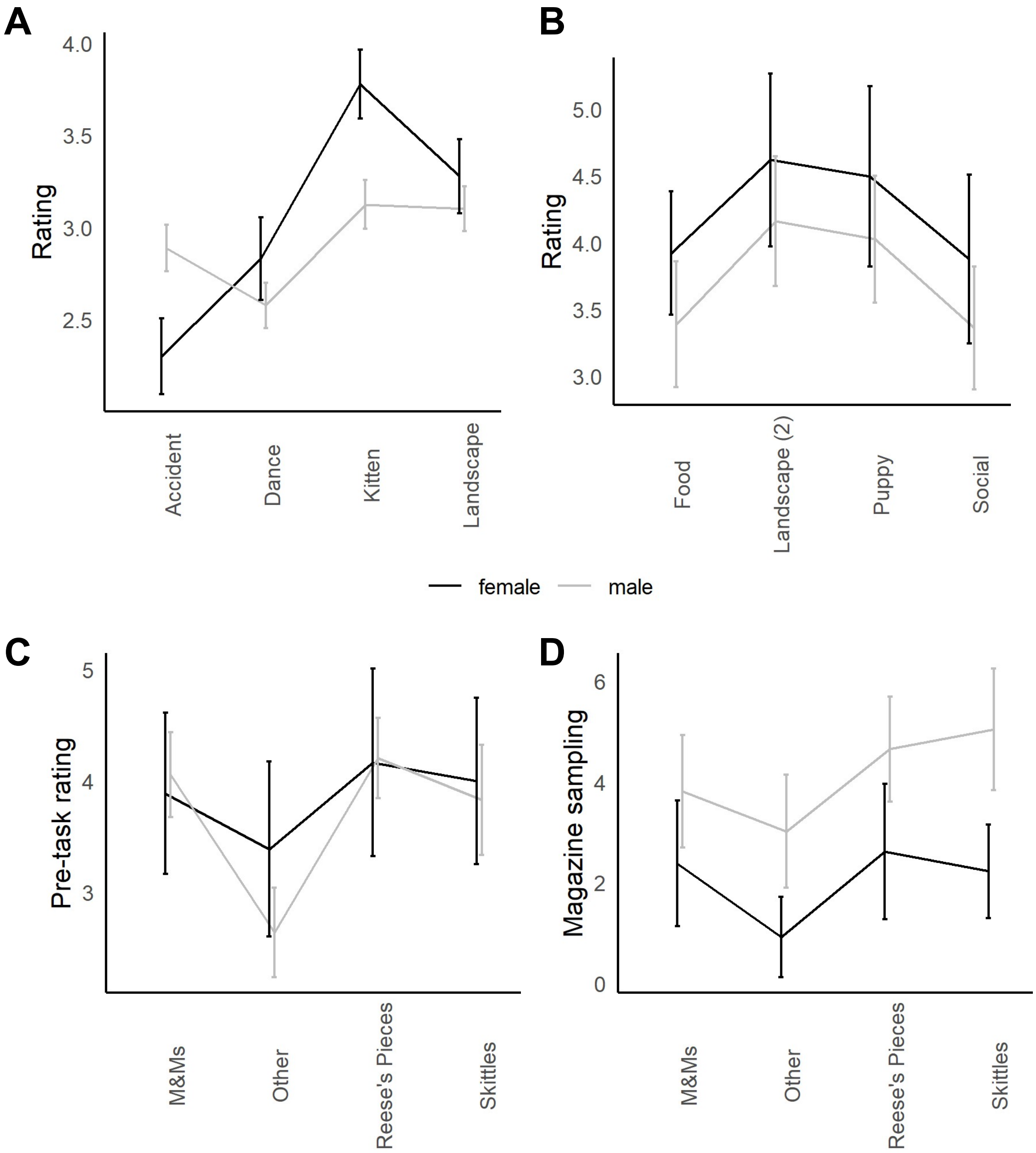

*Supplementary Figure 1.* Reward preferences by gender. A-B: Post-video ratings (1-5 stars) for the Movie Row task using the original Web-Surf (A) or a second (B) set of movies. C-D: Candy Row task. C: Pre-task ratings of enjoyment each reward (1-6). D: Number of magazine entries per trial for each reward type. Gender differences were only significant for the original Web-Surf videos (A), with males rating Accident videos more highly, and females rating Kitten videos more highly.

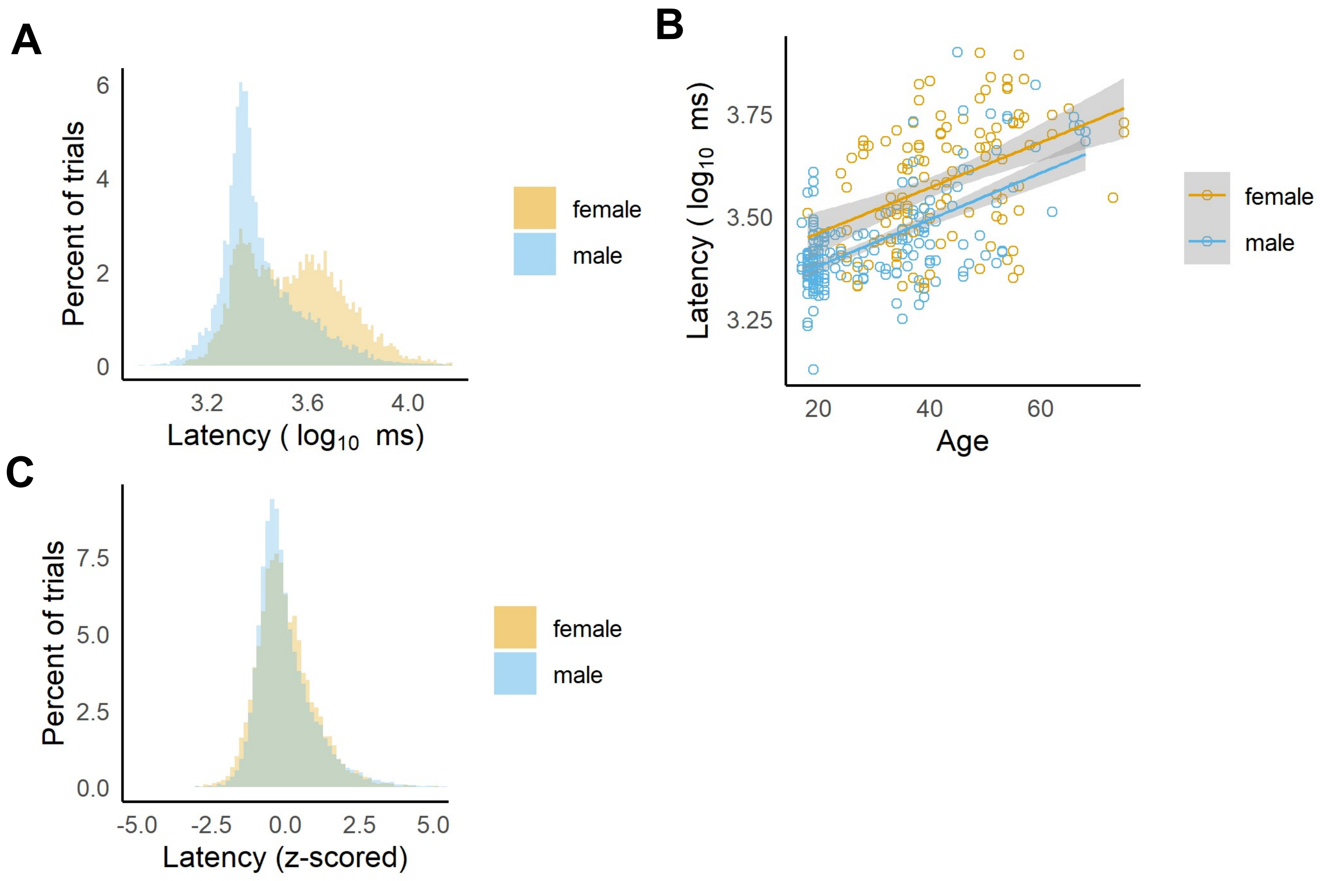

*Supplementary Figure 2.* Decision latencies. A: After $\log_{10}$ transformation, the distribution of decision latencies (across all trials from the first session for all participants) was bimodal for females and had a strong positive skew for males. B: Average decision latency ($\log_{10}$ milliseconds) was related to age and gender, with females having longer latencies on average, and latencies increasing with age. C: After z-scoring the $\log_{10}$ transformed decision latencies (within session), the differences between females and males was reduced, but males tended to have a stronger positive skew in their decision latencies.

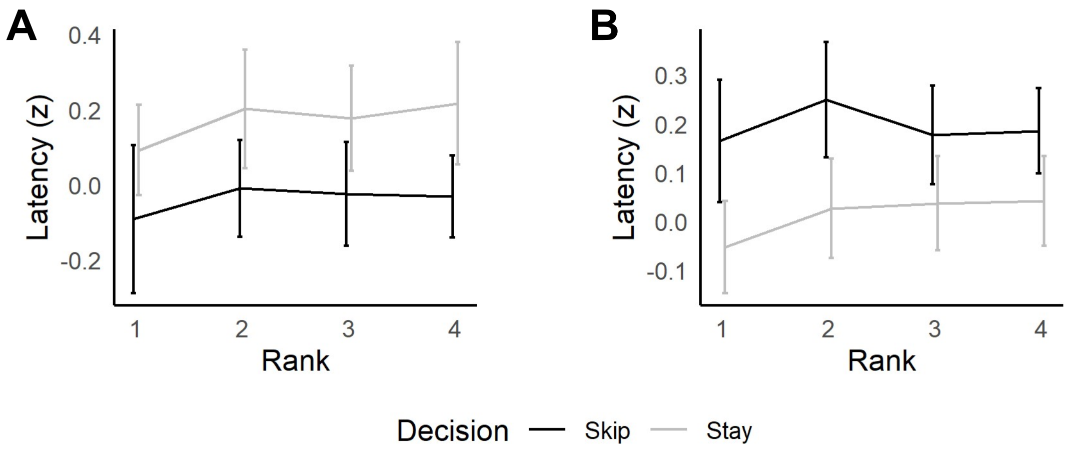

*Supplementary Figure 3.* Decision latency by post-task ranking and gender. Decision latencies were slower for stay decisions for females (A) and skip decisions for males (B). Overall, latencies were also faster for the favorite rewards (ranked 1). No significant interactions with rank were observed.

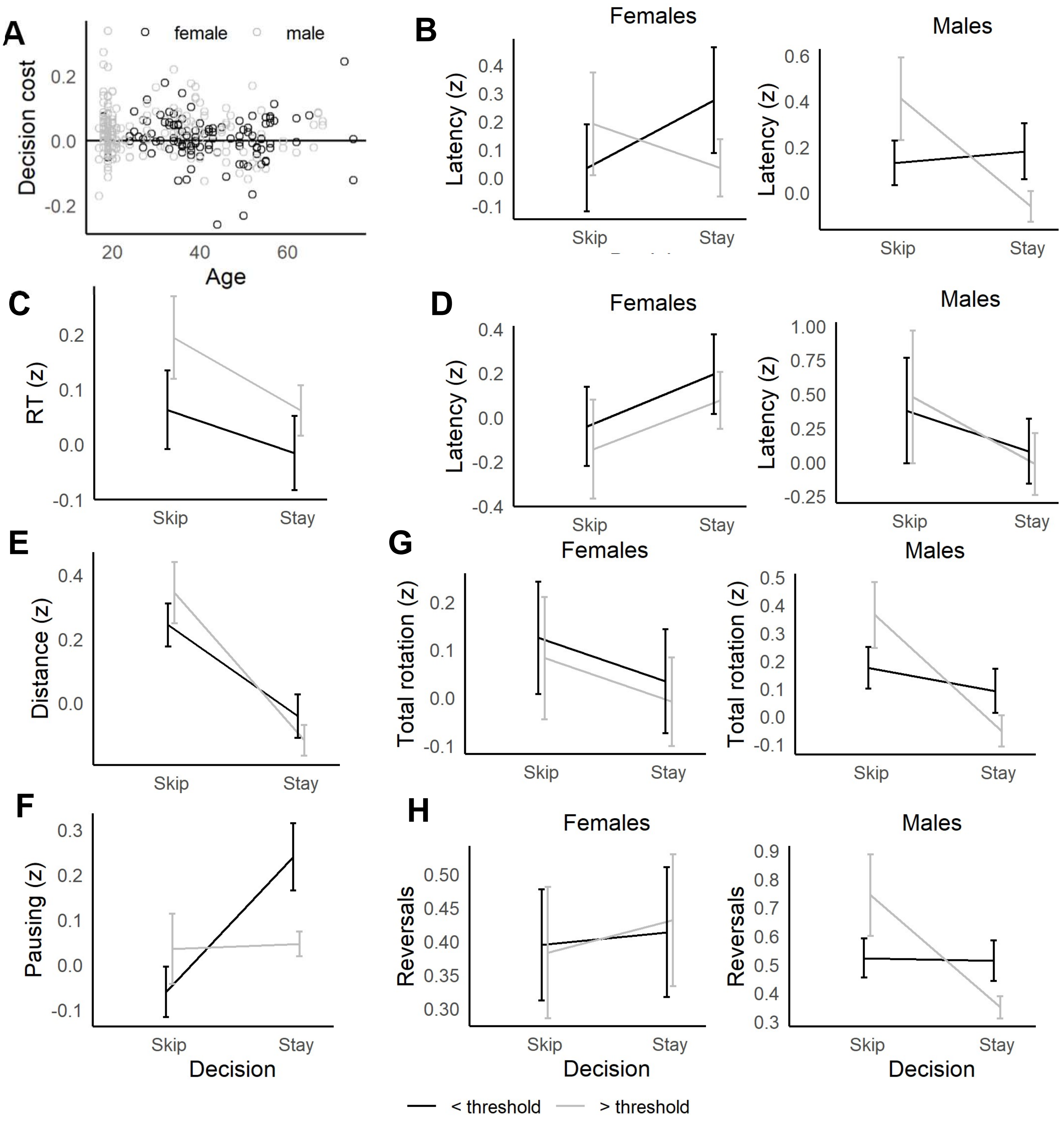

*Supplementary Figure 4.* Behavior when making choices inconsistent with delay thresholds. A: Average difference between inconsistent and consistent choices (Decision cost) versus age. Overall, 69% of males versus 56% of females had a positive decision cost (and had longer latencies when accepting offers above threshold versus below, or skipping offers below threshold versus above). This gender difference was related to age, with more participants 40 years old and younger having decision costs over 0 (71% of males, 59% of females, panel B) than those who were over 40 years of age (56% of males, 53% of females, panel D). C: Reaction times were longer when participants skipped good offers. E-H: Participants travelled farther (E) and paused longer (F) when accepting offers above threshold and skipping offers below threshold. G-H: A similar pattern was seen for the total amount of rotation (G) and number of reversals in rotation direction (H) in the offer zone, but only for males (right).

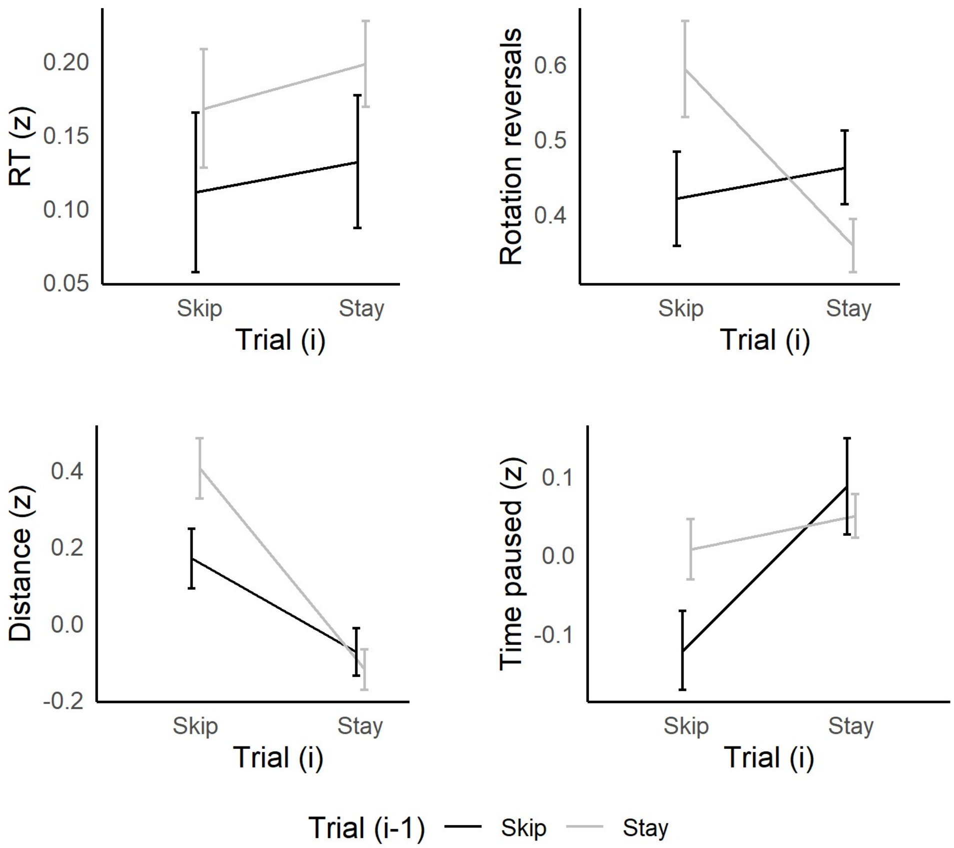

*Supplementary Figure 5*. Sequential choice behavior. Participants travelled farther, rotated farther and made more reversals in the direction of their rotation after previously skipping an offer (on trial i-1), compared to other sequences Skip/Skip sequences. These behaviors on Stay/Skip sequences were generally higher than other sequences (Stay/Stay, or Skip/Stay), except for pausing, which was more likely on average when participants accepted an offer. No significant differences were observed for reaction times.

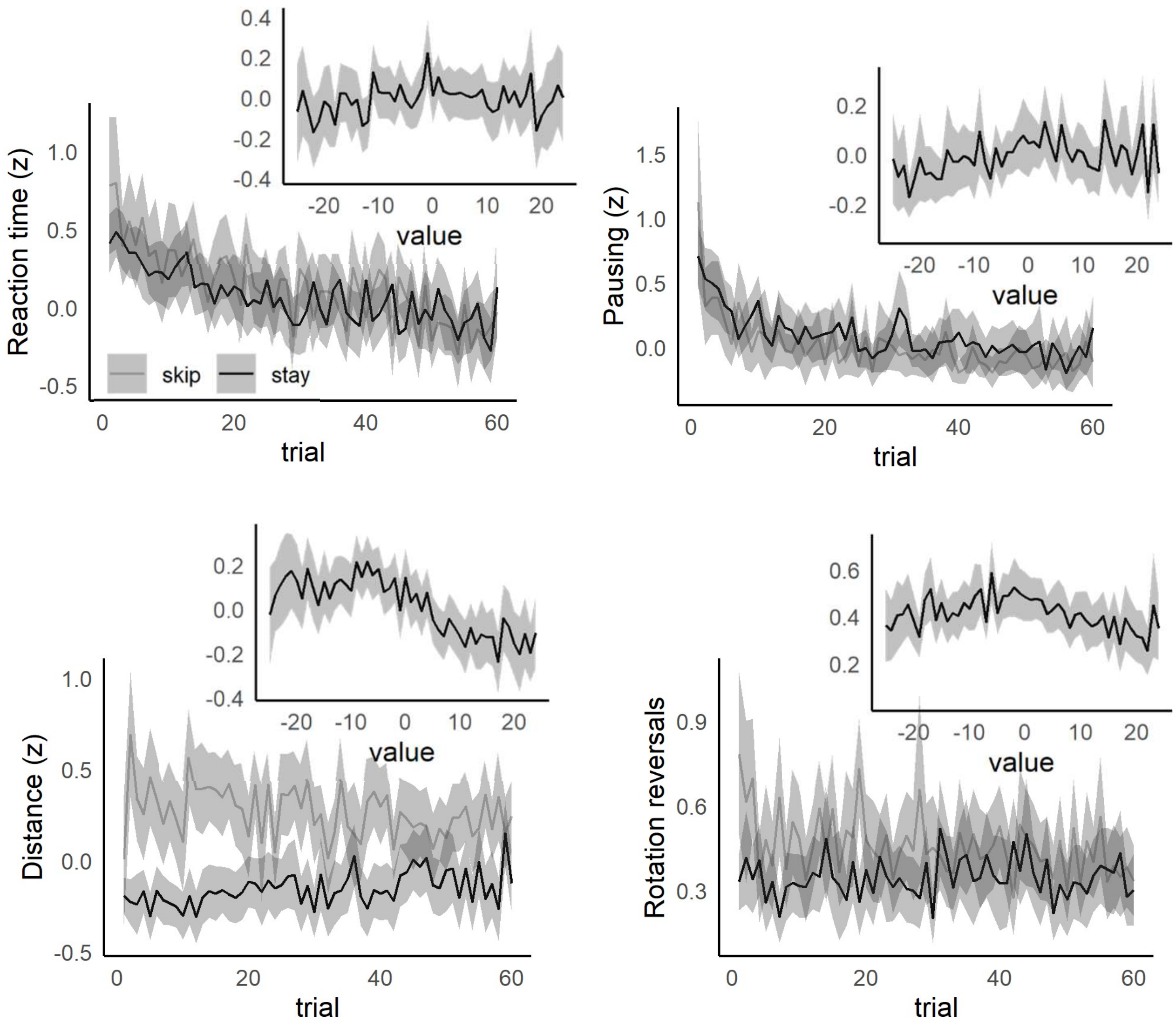

*Supplementary Figure 6*. Cross-session changes in behavior and relationship to deliberation. Across the session, reaction time and time spent paused decreased for both stay and skip decisions. For rotation reversals and distance travelled, a similar decrease across trials was observed for skip decisions. For each measure, plots relative to trial were calculated across participants, plots relative to value were calculated by binning values at 1 second intervals between -25 to +25 seconds, then averaging within participant before calculating means and confidence intervals. Lines indicate means, shaded areas indicate 95% confidence intervals.
